# Exposure duration modulates the response of Caribbean corals to global change stressors

**DOI:** 10.1101/2020.06.19.161711

**Authors:** HE Aichelman, CB Bove, KD Castillo, JM Boulton, AC Knowlton, OC Nieves, JB Ries, SW Davies

## Abstract

Global change is threatening coral reefs, with rising temperatures leading to repeat bleaching events (dysbiosis of coral hosts and their symbiotic algae) and ocean acidification reducing net coral calcification. Although global-scale mass bleaching events are revealing fine-scale patterns of coral resistance and resilience, traits that lead to persistence under environmental stress remain elusive. Here, we conducted a 95-day controlled-laboratory experiment to investigate how duration of exposure to ocean warming (28, 31°C), acidification (*p*CO_2_ = 400–2800 μatm), and their interaction influence the physiological responses of two Caribbean reef-building coral species (*Siderastrea siderea*, *Pseudodiploria strigosa*) from two reef zones of the Belize Mesoamerican Barrier Reef System. Every 30 days, calcification rate, total host protein and carbohydrate, chlorophyll *a* pigment concentration, and symbiont cell density were quantified for the same coral colony to characterize acclimatory responses of each genotype. Physiologies of the two species were differentially affected by these stressors, with exposure duration modulating responses. *Siderastrea siderea* was most affected by extreme *p*CO_2_ (~2800 μatm), which resulted in reduced calcification rate, symbiont density, and chlorophyll *a* concentration*. Siderastrea siderea* calcification rate initially declined under extreme *p*CO_2_ but recovered by the final time point, and overall demonstrated resistance to next-century *p*CO_2_ and temperature stress. In contrast, *P. strigosa* was more negatively impacted by elevated temperature (31°C). Reductions in *P. strigosa* calcification rate and total carbohydrates were consistently observed over time regardless of *p*CO_2_ treatment, with the greatest reductions observed under elevated temperature. However, nearshore colonies of *P. strigosa* maintained calcification rates under elevated temperature throughout all exposure durations, suggesting individuals from this environment may be locally adapted to the warmer temperatures characterizing their natal reef zone. This experiment highlights how tracking individual coral colony physiology across broad exposure durations can capture acclimatory responses of corals to global change stressors.

## 1. Introduction

Since the Industrial Revolution, anthropogenic activities have increased the partial pressure of atmospheric carbon dioxide (*p*CO_2_), which has warmed the atmosphere by approximately 0.6°C (Pörtner et al., 2019). As atmospheric temperatures increase, so do sea surface temperatures (SSTs), a trend that has continued since 1970 (Pörtner et al., 2019). Increasing atmospheric *p*CO_2_ has also caused surface ocean pH to decrease at a rate of 0.017 to 0.027 units per decade since the 1980s, potentially exacerbating the impacts of warming SSTs on marine organisms (e.g. Pörtner et al., 2019). The resulting warming and acidification have impacted organisms across the globe, as thermal niches shift and habitats rapidly change (Morley et al., 2018; Parmesan and Yohe, 2003; Pörtner et al., 2019). The negative effects of global change are predicted to strengthen and, under the Intergovernmental Panel on Climate Change’s (IPCC) most extreme emissions scenario (RCP8.5), oceans are expected to take up 5 to 7 times more heat and decrease by 0.3 pH units by 2100 (Pörtner et al., 2019).

Coral reefs are valuable economic and ecological resources (Costanza et al., 2014) that are particularly vulnerable to ocean warming and acidification. The high biodiversity of coral reefs depends on the obligate symbiosis between the coral animal and its symbiotic algae of the family Symbiodiniaceae (previously genus *Symbiodinium*; LaJeunesse et al., 2018). This symbiosis is sensitive to thermal anomalies and, because tropical coral species live within 1°C of their upper thermal limit, even slight increases in SST can result in bleaching (breakdown of the coral-algal symbiosis) and ultimately mortality if the symbionts fail to repopulate the coral host. Global coral bleaching events are occurring with increasing frequency and severity as SST continues to increase (Hughes et al., 2017).

Ocean acidification is the result of increased atmospheric CO_2_ dissolving into seawater, reducing its pH, and altering its carbonate chemistry (Doney et al., 2009; Orr et al., 2005). This alteration of carbonate chemistry includes a decrease in carbonate ion concentration ([CO_3_^2−^]), which reduces the saturation state of seawater with respect to aragonite (Ω_narag_)—which can make it more challenging for corals to build their aragonite skeletons (Doney et al., 2009; Kleypas et al., 1999). Laboratory experiments have shown that ocean acidification conditions projected for the end of next century can have negative (Comeau et al., 2013; Hoegh-Guldberg et al., 2007; Horvath et al., 2016; Kroeker et al., 2010), neutral (Reynaud et al., 2003; Ries et al., 2010), and parabolic (Castillo et al., 2014) impacts on coral calcification, while field experiments have yielded more negative outcomes (Albright et al., 2018; Comeau et al., 2019; Jokiel et al., 2008; Kline et al., 2019). The direction and magnitude of calcification responses to acidification are influenced by many factors (Kornder et al., 2018), including ability to regulate calcifying fluid chemistry in support of calcification (e.g. Ries, 2011), species differences (Bove et al., 2019; Okazaki et al., 2017), calcification rate under ambient conditions (Shaw et al., 2016), CO_2_-induced fertilization of photosynthesis (Castillo et al., 2014), coral gender (Holcomb et al., 2012), experimental duration (Kline et al., 2019), co-occurring thermal stress (Anthony et al., 2011; Kroeker et al., 2013), and heterotrophic capacity (Cohen and Holcomb, 2009; Towle et al., 2015). Coral calcification in response to temperature stress is similarly complicated by a number of factors, and a meta-analysis by Kornder et al. (2018) highlighted strong taxonomic variation and that thermal stress is more pronounced in adults and during the summer. Similar to acidification, coral calcification under elevated temperature can be mediated by heterotrophic capacity (Aichelman et al., 2016; Grottoli et al., 2006; Rodolfo-Metalpa et al., 2008; Towle et al., 2015). In addition to considering calcification rate, energetic reserves are critical to coral health and resistance to stressors, and have been associated with bleaching susceptibility (Anthony et al., 2009; Grottoli et al., 2014; Levas et al., 2018) and in determining whether a bleaching event will lead to mortality (Anthony et al., 2009; Grottoli et al., 2006).

Fewer studies consider the combined effects of temperature and acidification stress and, similar to studies investigating the effects of independent stressors, such experiments have produced variable results. Several studies have demonstrated that elevated temperatures have stronger negative effects on the coral holobiont (combination of coral animal, algal symbiont, and associated microbiota) when compared to acidification, including negative effects on calcification rates (Anderson et al., 2019; Schoepf et al., 2013), larval development (Chua et al., 2013), and survivorship (Anderson et al., 2019). Although no studies have found synergistic effects of temperature and acidification on coral calcification (i.e. combined effects are greater than sum of individual effects), numerous studies have demonstrated that the effects of temperature and acidification stress can be additive, in terms of impacts on coral calcification (Agostini et al., 2013; Edmunds et al., 2012; Horvath et al., 2016; Kornder et al., 2018; Prada et al., 2017; Rodolfo-Metalpa et al., 2011) and metabolism (Agostini et al., 2013). A more complete understanding of the combined effects of global change stressors will require investigations of multiple species and stressors across longer timescales with a focus on multiple physiology metrics.

The effects of global change on the coral holobiont not only vary by the stressors in question, but also by species. Species differences in response to global change stressors have been observed in coral calcification rate (Bove et al., 2019; Edmunds et al., 2012; Edmunds et al., 2019; Okazaki et al., 2017) and recovery of energetic reserves through time (i.e. total soluble lipid; Levas et al., 2018). Additionally, spatial scale appears to play a role in some corals’ responses to global change, with differential stress tolerance observed across populations along a reef system (e.g. the Great Barrier Reef; Dixon et al., 2015) and across reef zones (Castillo et al., 2012; Kenkel et al., 2013a; Kenkel et al., 2013b; Kenkel and Matz, 2016). Differences in coral thermal tolerance can occur on spatial scales as small as between tidal pools in American Samoa (Bay and Palumbi, 2014; Oliver and Palumbi, 2011; Palumbi et al., 2014), illustrating that adaptation and/or acclimation to fine scale environmental differences can play a role in determining coral response to global change stressors.

In addition to understanding stress responses across species and spatial scales, considering how duration of stress exposure affects the physiological response of the coral holobiont is critical (McLachlan et al., 2020). This goal is complicated by the difficulty of executing long-term laboratory (*ex situ*) experiments, and it is therefore relatively rare for studies to track coral physiology under controlled conditions for extended periods of time. However, several studies have been conducted for approximately 90 days or more (e.g., Anderson et al., 2019; Bove et al., 2019; Castillo et al., 2014; Comeau et al., 2019; Kline et al., 2019), and each reveal nuanced patterns of stress and resilience in corals. For example, Comeau et al. (2019) observed that acidification (1,050 μatm *p*CO_2_) caused a rapid, but species-specific, alteration of calcifying fluid chemistry in four coral and two calcifying algae species throughout the entire one-year duration of the experiment (Comeau et al., 2019). Additionally, by measuring *S. siderea* growth every 30 days throughout a 90-day experiment, Castillo et al. (2014) showed that calcification responses to acidification (604 μatm *p*CO_2_) varied substantially through time—with calcification rates increasing in response to moderate acidification (604 μatm *p*CO_2_) between 0 and 60 days, and decreasing between 60 and 90 days. Finally, Levas et al. (2018) tracked corals for 11 months following experimental bleaching and found interspecific differences in the timing of recovery. Specifically, *Porites divaricata* initially catabolized lipids and decreased calcification but largely recovered within 11 months, while *P. astreoides* fully recovered within 1.5 months after increasing feeding and symbiont nitrogen uptake (Levas et al., 2018). It is therefore clear that tracking coral physiology through time can provide valuable insights into how corals respond to short-, moderate-, and long-term global change stress.

Here, two ecologically important reef-building coral species (*Siderastrea siderea* and *Pseudodiploria strigosa*) from two reef zones (forereef and nearshore) of the Belize Mesoamerican Barrier Reef System (MBRS) were maintained under a fully crossed acidification (ca. 400 μatm [present day], ca. 640 μatm [next century], ca. 2800 μatm [extreme]) and temperature (28, 31°C) experiment for 95 days. In order to characterize the responses of these species to projected global change, holobiont physiology of each coral colony was monitored every 30 days (exposure duration: 0-30 days = T_0_-T_30_ = short-term, 30-60 days = T_30_-T_60_ = moderate-term, 60-95 days = T_60_-T_95_ = long-term), including metrics for both the coral host (calcification rate, total protein, total carbohydrates) and symbiont (symbiont cell density, chlorophyll *a* pigment concentration). The work presented here elucidates the impact of exposure duration on corals’ acclimatory response to global change stressors.

## 2. Materials and methods

### (2.1) Coral collection and experimental design

Data presented here are from an experiment run in parallel with an experiment published by Bove et al. (2019). Therefore, experimental design and culturing conditions are similar to those presented therein. However, data presented here pertain to a different suite of coral individuals than explored in Bove et al. (2019), and only two species (instead of four) are explored in the present study. Timing of the two experiments is also staggered by 30 days; for comparison time T_0_ in the present experiment corresponds to the “pre-acclimation period” in Bove et al. (2019). Methods specific to this experiment are presented below. However, readers are referred to Bove et al. (2019) for methods that pertain to both studies, such as those relating to the control and monitoring of seawater carbonate chemistry parameters.

Three colonies of *Siderastrea siderea* and *Pseudodiploria strigosa* were collected from a nearshore (NS; Port Honduras Marine Reserve, PHMR; 16°11’23.5314”N, 88°34’21.9360”W) and a forereef (FR; Sapodilla Cayes Marine Reserve, SCMR; 16°07’00.0114”N, 88°15’41.1834”W) site along the southern portion of the Belize Mesoamerican Barrier Reef System (MBRS) in June 2015 (N=3/species/site; Figure 1). Colonies were separated by at least 5 m to maximize the likelihood of obtaining genetically distinct individuals. Following collection, all coral colonies (3 colonies x 2 reef zones x 2 species = 12 putative genotypes) were transported to the Northeastern University (NU) Marine Science Center and fragmented into 24 genetically identical fragments. One FR *P. strigosa* colony did not survive fragmentation, leaving a total of 3 genotypes for NS and FR *S. siderea,* 3 genotypes for NS *P. strigosa*, and 2 genotypes for FR *P. strigosa* genotypes (total of 6 *S. siderea* and 5 *P. strigosa* genotypes). After fragmentation, coral fragments were allowed to recover for 23 days in natural flow-through seawater with salinity and temperature (±SD) of 30.7±0.8 and 28.2±0.5°C, respectively. After recovery, temperature and *p*CO_2_ were incrementally adjusted over a period of 20 days until target treatment conditions were achieved. During this time, temperatures of the elevated temperature treatments were increased by 0.4°C every 3 days and *p*CO_2_ was adjusted by 0 μatm (present day), +30 μatm (end of century), and +240 μatm (extreme) every 3 days. The six experimental treatments consisted of a full factorial design of two temperatures (target 28, 31°C) and three *p*CO_2_ levels (target 400, 700, 2800 μatm). Coral fragments were distributed such that four replicate fragments from each genotype were represented in each of the six experimental treatments. The six experimental treatments were replicated in three aquaria, for a total of 18 42L acrylic aquaria. Each aquaria was illuminated on a 10:14 h light:dark cycle with 300 μmol photons m^−2^ s^−1^ of photosynthetically active radiation (PAR). The experimental system used natural flow-through seawater and coral fragments were fed every other day with a mixture of frozen adult *Artemia* sp. and freshly hatched *Artemia* sp.

**Figure 1.**
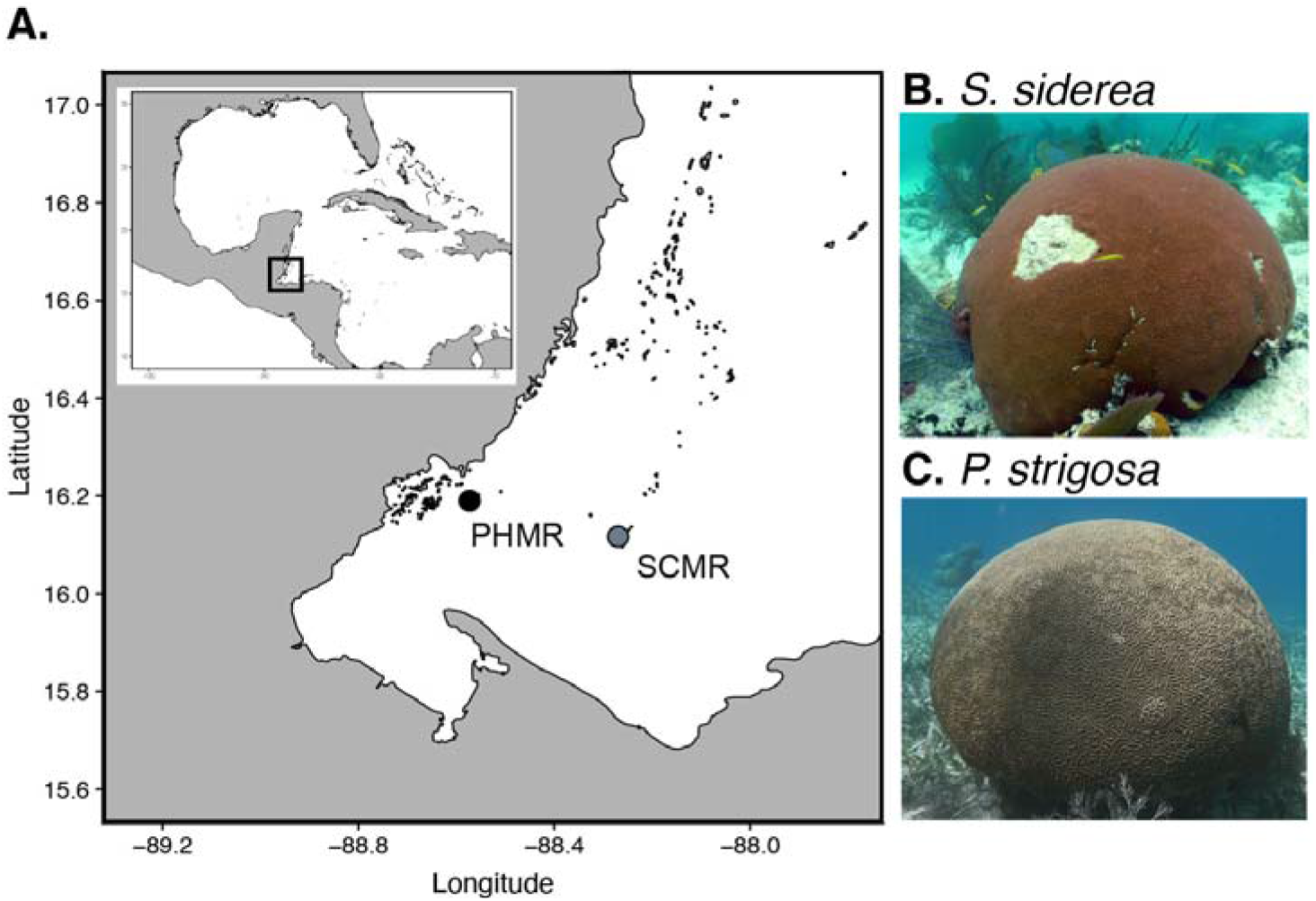
**(A)** Map of coral colony collection sites from the forereef (SCMR = Sapodilla Cayes Marine Reserve) and nearshore (PHMR = Port Honduras Marine Reserve) sites of the Belize Mesoamerican Barrier Reef System (MBRS). **(B)** Example of *Siderastrea siderea* colony (photo credit: K.D. Castillo). **(C)** Example of *Pseudodiploria strigosa* colony (photo credit: H.E. Aichelman).

Coral fragments were maintained in treatment conditions for a total of 95 days (9 August 2015 – 12 November 2015). Average (±SE) *p*CO_2_ and temperature conditions throughout the experimental period were: 358 (±131) μatm, 28.2 (±0.1) °C; 424 (±40) μatm, 31.3 (±0.1) °C; 674 (±21) μatm, 28.0 (±0.1) °C; 606 (±36) μatm, 30.9 (±0.1) °C; 2750 (±161) μatm, 28.4 (±0.1) °C; 2917 (±174) μatm, 31.1 (±0.1) °C. Seawater parameters (including temperature, *p*CO_2_, salinity) at each time point are shown in Figure S1 and all measured and calculated seawater parameters are reported in Tables S1 and S2, respectively.

Approximately every 30 days (exposure duration: T_0_-T_30_ days = short-term, T_30_-T_60_ days = moderate-term, T_60_-T_95_ days = long-term), a fragment of each coral colony was removed from each of the six experimental conditions, flash frozen in liquid nitrogen, and stored at −80°C for subsequent analysis aimed at tracking genet-level physiology through time. In the event of mortality that yielded insufficient coral fragments for sampling at all time points, corals were preferentially sampled at the end of the experiment (long-term exposure: T_95_) instead of after moderate-term exposure (T_60_). As a result, sample sizes are lower for both species at T_60_ compared to the other time points. Following completion of the experiment, all remaining coral fragments were flash frozen in liquid nitrogen and maintained at −80°C until subsequent processing, at which time fragments were airbrushed to remove host tissue and symbiont cells. Tissue slurries were homogenized using a Tissue-Tearor (Dremel; Racine, WI, USA) and centrifuged to pelletize the symbiont cells. Coral host tissue and symbiont fractions were separated for subsequent physiological assays.

### (2.2) Coral host physiology measurements

Coral growth rates were estimated over the course of the experiment using the buoyant weight technique (Davies, 1989). Buoyant weight measurements were obtained in triplicate for each coral fragment at each of the four time points (T_0_, T_30_, T_60_, T_95_), averaged, and normalized to the coral specimens’ surface area (see methods below). As in Bove et al. (2019), a subset of fragments from both species were used to confirm the relationship between buoyant weight and dry weight. These two measurements were correlated for both species (*S. siderea* R^2^ = 0.90, p < 0.001; *P. strigosa* R^2^ = 0.81, p < 0.001), indicating that change in buoyant weight should reflect a proportionate change in dry weight. Equations used to calculate dry weight from buoyant weight are shown below. Dry weight was converted from g to mg, corrected to surface area of each fragment and to number of days in experimental treatment to calculate calcification rate (mg cm^−2^ day^−1^).

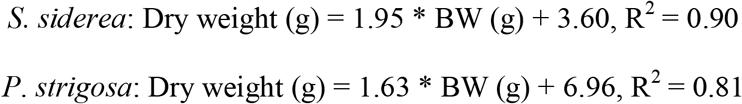

Growing surface area was quantified in triplicate from observed live tissue in photos of coral fragments taken at each timepoint using ImageJ software (Rueden et al., 2017). The same surface area values of each coral fragment were used to normalize all host and symbiont physiology parameters within an experimental timepoint.

Because corals were frozen on the same day for each time point, there was no need to correct for number of days in experimental treatment for physiological metrics other than calcification rate. Total coral host protein content was quantified from host tissue slurry using a bicinchoninic acid (BCA) protein assay. Host tissue slurry was vortexed with glass beads for 15 minutes and then centrifuged for 3 minutes at 4000 RPM. Next, 15 μL of the centrifuged sample was added to 235 μL artificial seawater along with 250 μL of Bradford reagent. After samples were mixed, absorbance was measured in a BioSpectrometer (Eppendorf, Hauppauge, NY, USA) at 562 nm. Coral protein concentrations were calculated using a standard curve of bovine serum albumin ranging from 0 to 1000 μg mL^−1^ and normalized to living coral surface area.

Total host carbohydrates were quantified using the phenol-sulfuric acid method (as in Masuko et al., 2005), which measures all monosaccharides, including glucose—the major photosynthate translocated from symbiont to coral host (Burriesci et al., 2012). An aliquot of coral host tissue was diluted to 50 μL with artificial seawater (Instant Ocean Sea Salt), to which 150 μL of sulfuric acid and 30 μL of 5% phenol were added. Following a 5-minute incubation at 90°C and another 5-minute incubation at room temperature, absorbance at 490 nm was measured in a spectrophotometer (Synergy H1 Microplate Reader; BioTek Instruments; VT, USA). Carbohydrate concentrations were calculated using a standard curve of D-glucose solutions ranging from 0.039 to 2 mg mL^−1^ and normalized to living coral surface area.

### (2.3) Symbiodiniaceae physiology measurements

Symbiont cells were quantified using the hemocytometer method similar to Rodrigues and Grottoli (2007). After vortexing the symbiont pellet, a 1:1 Lugol’s iodine and formalin solution was added for contrast and cell preservation. Triplicate 10 μL subsamples were counted on a hemocytometer using a light microscope, averaged, and normalized to slurry volume and live coral tissue surface area.

Symbiont photosynthetic pigments (chlorophyll *a,* abbreviated Chl *a*) were quantified spectrophotometrically following the method of Marchetti et al. (2012). Briefly, 40 mL of 90% acetone was added to the symbiont pellet, homogenized, then stored in the dark for 24 hours. 100 μL of each sample was then diluted in 7.9 mL of 90% acetone. A 10AU Field and Laboratory Fluorometer (Turner Designs, San Jose, CA) was used to measure the initial concentration (R_b_), then 2 drops of 10% HCl was added to the sample tube, after which a second fluorometer reading was taken (R_a_). Total Chl *a* content (μg L^−1^) was calculated using the equation below, where 0.548 is a calibration constant specific to the fluorometer used, 40 mL is the volume of acetone left overnight, and 80 is the dilution factor. Total Chl *a* was then normalized to live coral surface area to get units of μg Chl *a* cm^−2^.

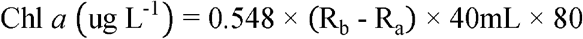

### (2.4) Statistical analyses

All statistical analyses were completed using R version 3.5.2 (R Core Team, 2017). A series of linear models (package *lmer*) were fitted for each species and each individual physiology parameter using a forward model selection method. The best fit model was derived by starting with the intercept-only model and then using forward-selection to incorporate additional parameters, starting with the most significant parameter, until further addition of parameters did not significantly improve the model fit. Additional parameters were retained in the model if they were significant (p < 0.05) and produced smaller AIC values (Akaike, 1974). These parameters included fixed effects of time, temperature, *p*CO_2_, and reef zone, which were all coded as factors. Parameter interactions were only considered if those two parameters were already significant and included in the model. A random effect of genotype was included in all models to account for physiological variation across genotypes. Note that for calcification rate data, multiple fragments of each genotype were represented at each time point. Because genotype is included in the model as a random effect, multiple fragment numbers do not artificially increase the sample size and instead only increase the precision of the rate measurement for that colony. The linear models used for each individual physiology parameter are included along with summary statistics for *S. siderea* and *P. strigosa* in Table S3. Post-hoc pairwise comparisons of significant main effects were assessed using a Tukey’s HSD test, implemented in the *lsmeans* function with the option “adjust = tukey”. Summary statistics for all post-hoc comparisons are reported in Table S4.

A Principal Components Analysis (PCA) was constructed using the *FactoMineR* package (Lê et al., 2008) to assess how overall physiologies were modulated through time for each species. All physiology parameters were log-transformed, and calcification rates were x+2 log-transformed. Only individual coral fragments for which all physiology parameters were present (calcification rate, total protein, total carbohydrate, symbiont density, Chl *a*) were included in this analysis. Significance of each factor in the PCA was assessed using the *adonis()* function in the *vegan* package (Oksanen, 2011), which was run with 10,000 permutations using the model below. Summary statistics for all *Adonis* tests are reported in Table S5.

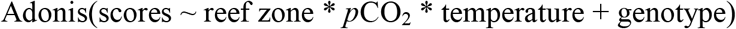

Correlation matrices of all host and symbiont physiology parameters for both species through time were built using the *corrplot()* function with a significance threshold of p = 0.05. The impacts of temperature and *p*CO_2_ on all host and symbiont physiology parameters of only *P. strigosa* were assessed via linear regression modeling, as no noteworthy correlations were found for *S. siderea*. To estimate significance of the predictors and their interactions, increasingly parsimonious, nested linear models (using *lmer()*) were compared with likelihood ratio tests. Conditional R-squared values (accounting for both fixed and random effects) of the regressions were determined using the *r.squaredGLMM()* function in the *MuMIn* package. Summary statistics for all linear regressions are reported in Table S6.

All data and code used for the analyses presented herein can be found on the GitHub repository associated with this publication (https://github.com/hannahaichelman/TimeCourse_Physiology).

## 3. Results

### (3.1.) Effects of thermal and acidification stress on *S. siderea* and *P. strigosa* calcification rates

Experiment duration (p < 0.001), *p*CO_2_ (p < 0.001), and the interaction of experiment duration and *p*CO_2_ (p = 0.007), significantly influenced calcification rates of *S. siderea* (Figure 2A). Under extreme *p*CO_2_ conditions, *S. siderea* calcification rates were significantly reduced relative to both present day (Tukey p < 0.001) and next century (Tukey p = 0.017) *p*CO_2_ treatments. Through time, calcification rates of *S. siderea* in extreme *p*CO_2_ conditions declined between T_30_ and T_60_ (Tukey p < 0.001), and also over the entire duration of the experiment (T_30_ to T_95_; Tukey p < 0.001). Temperature was not included in the best-fit model, and therefore did not have a significant effect on *S. siderea* calcification.

*P. strigosa* calcification rates were significantly affected by experiment duration (p < 0.001), *p*CO_2_ (p < 0.001), temperature (p < 0.001) and reef zone (p = 0.036) (Figure 2B). Calcification rates were reduced at elevated temperature (31°C) relative to control temperature (Tukey p < 0.001) and nearshore corals calcified faster than forereef corals (Tukey p = 0.039). When compared to the present day *p*CO_2_ treatment, calcification rates were also reduced under extreme (Tukey p < 0.001) and next century *p*CO_2_ conditions (Tukey p = 0.017). A significant interaction of temperature and experimental duration was also detected for *P. strigosa* calcification rates (p < 0.001), with *P. strigosa* calcification rates decreasing between T_30_ and T_60_ under both control and elevated temperatures (Tukey p < 0.05); however, these reductions were no longer detectable after moderate- and long-term exposure (T_60_ and T_95_). When considering the full duration of the experiment (T_0_ to T_95_), *P. strigosa* calcification rates decreased under elevated temperature, but not under control temperature (Figure 2B). Additionally, a significant interaction between reef zone and temperature on *P. strigosa* calcification rate was detected (p = 0.031), with elevated temperatures more negatively influencing calcification of forereef corals than nearshore corals (Tukey p = 0.024). Lastly, the interaction between temperature and *p*CO_2_ significantly affected *P. strigosa* calcification rates (p = 0.0009). There were no significant differences in *P. strigosa* calcification rates amongst *p*CO_2_ treatments under elevated temperature treatments; however, calcification rates in control temperatures were significantly reduced under extreme *p*CO_2_ compared to the present day *p*CO_2_ conditions (Tukey p < 0.001).

**Figure 2.**
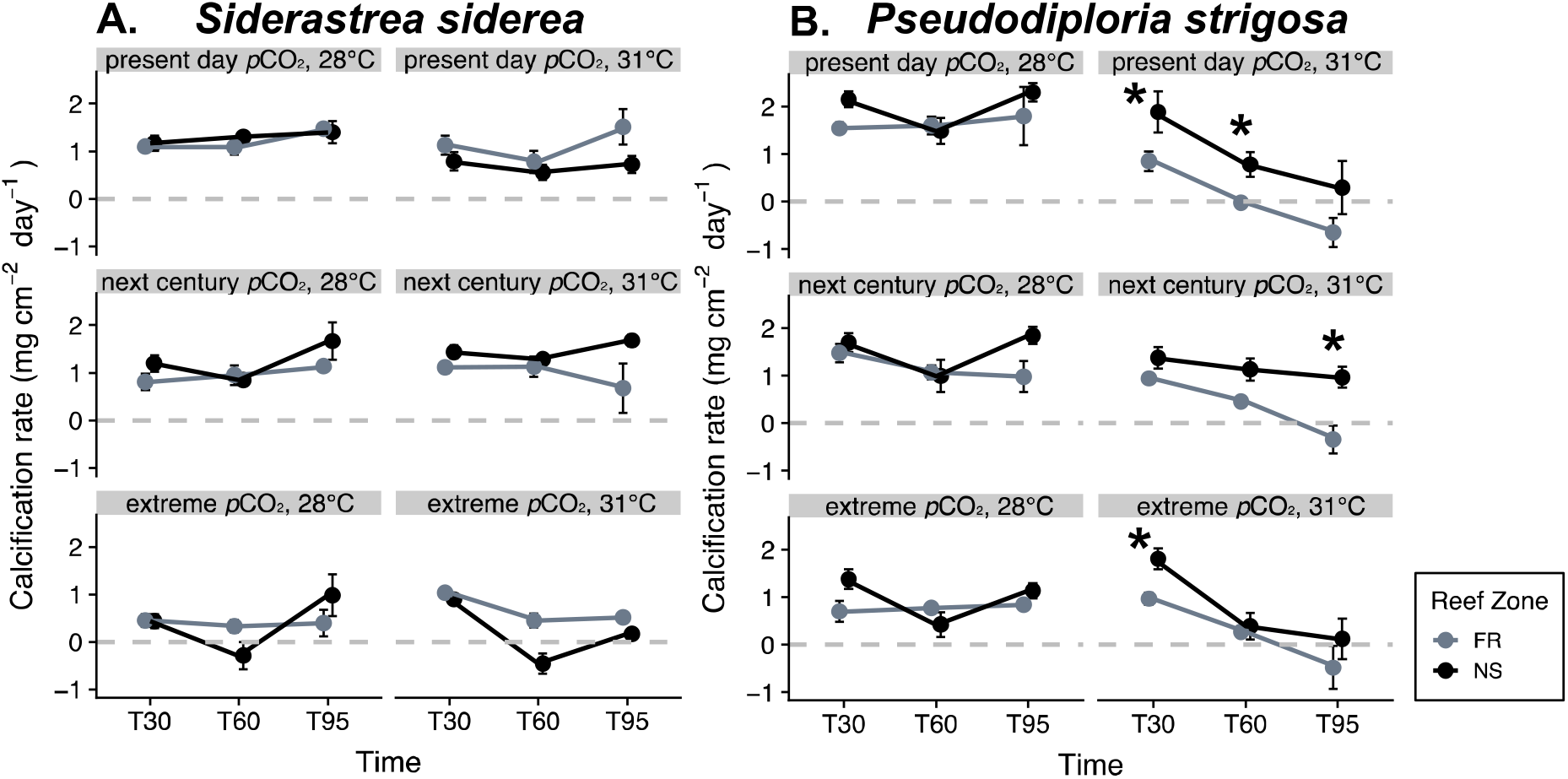
*Siderastrea siderea* **(A)** and *Pseudodiploria strigosa* **(B)** calcification rate (mg cm^−2^ day^−1^) at each experimental time point (short-term = T_30_; moderate-term = T_60_; long-term = T_95_). Facets represent each of the six treatments (*p*CO_2_: present day [~400 μatm], next century [~640 μatm], extreme [~2800 μatm]; temperature: 28°C, 31°C) and, within a facet, data are separated by reef zone (“FR” = forereef; “NS” = nearshore). Points represent mean calcification rates since the previous time point (i.e. T_30_ represents calcification between T_0_ and T_30_). Asterisks (*) indicate significant (Tukey p < 0.05) differences in calcification rates between reef zones within a time point. Error bars represent standard error. For *S. siderea* (A), each data point represents three colonies. For *P. strigosa* (B), each forereef point represents 2 colonies and each nearshore point represents 3 colonies, except at the extreme *p*CO_2_/28°C treatment at T_60_ and T_95_, where only one forereef colony is represented due to mortality.

### (3.2) Effects of thermal and acidification stress on *S. siderea* and *P. strigosa* host energy reserves

#### (3.2.1) *Siderastrea siderea* total protein and total carbohydrates

Both experiment duration (p < 0.001) and temperature (p < 0.01) had significant effects on *S. siderea* total protein content (Figure 3A), with elevated temperatures reducing total protein concentrations relative to corals in control temperatures (Tukey p = 0.01). Regardless of *p*CO_2_ and temperature treatments, *S. siderea* total protein content increased through time, with T_95_ exhibiting higher mean protein concentrations than T_0_ (Tukey p = 0.03). Temperature was the only factor that influenced total carbohydrates of *S. siderea* (Figure 3C; p < 0.01), with significantly less carbohydrate at elevated temperature relative to the control temperature treatment (Tukey p = 0.003).

**Figure 3.**
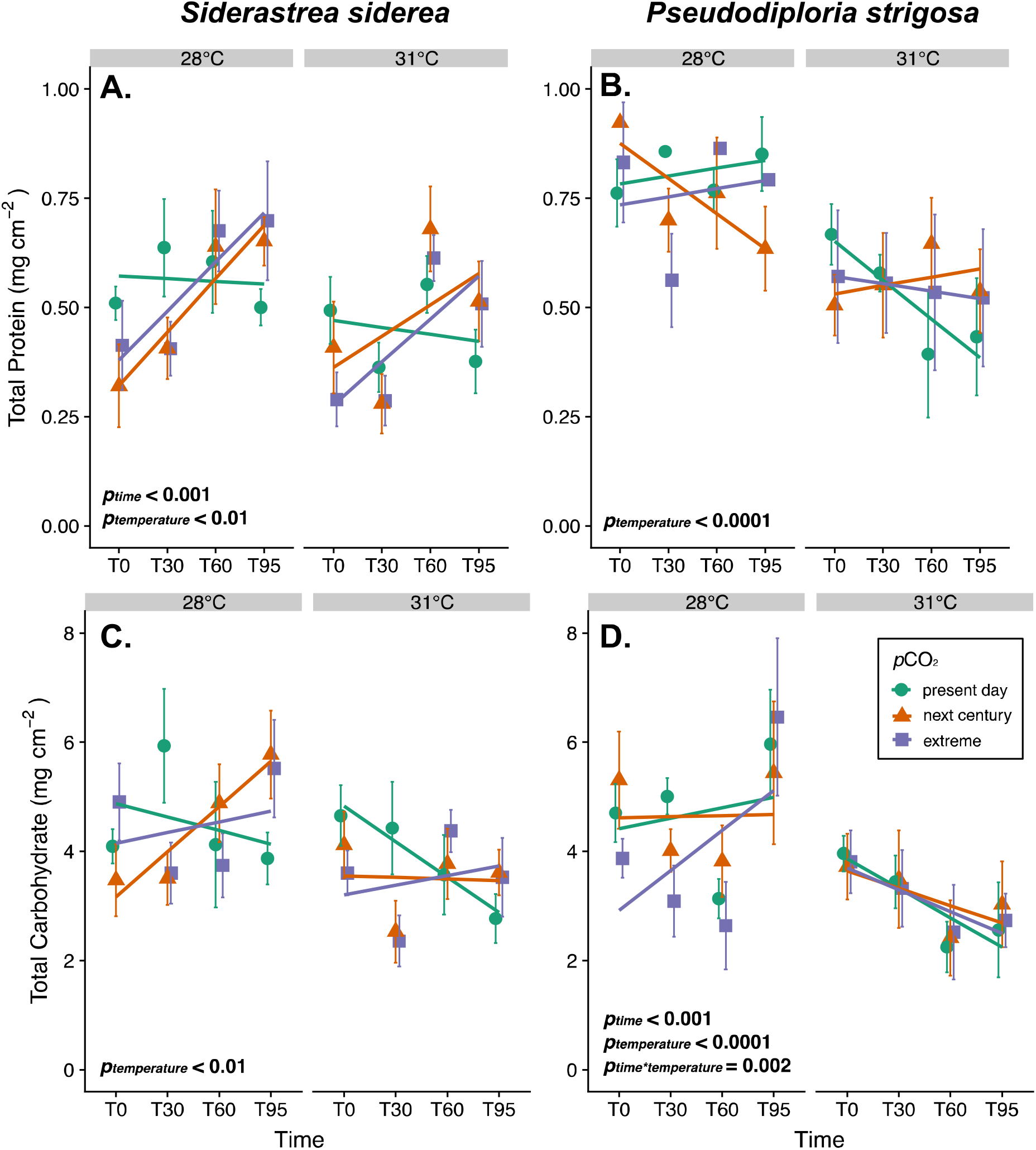
Host energy reserves (total protein **[A,B]** and total carbohydrate **[C,D]**) of *Siderastrea siderea* **(A,C)** and *Pseudodiploria strigosa* **(B,D)** across four experimental durations (day 0 = T_0_; day 30 [short-term] = T_30_; day 60 [moderate-term] = T_60_; day 95 [long-term] = T_95_). Within each panel, results are faceted by temperature treatment (28, 31°C) and colored by *p*CO_2_ treatment (present day [~400 μatm] = green, next century [~640 μatm] = orange, extreme [~2800 μatm] = purple). Each point is an average of corals from nearshore and forereef reef zones with n = 4-6 (*S. siderea*) and n = 3-6 (*P. strigosa*) distinct fragments (n = 1 / genotype). Significant factors are indicated in each panel. Lines represent linear fits (using ggplot2 *stat_smooth()* method to visualize differences regardless of model) for each treatment through time, and error bars represent standard error at each time point.

#### (3.2.2) *Pseudodiploria strigosa* total protein and total carbohydrates

For *P. strigosa,* temperature was the only factor that significantly influenced total host protein (p < 0.001), with reduced protein concentrations under elevated temperatures compared to control temperature treatments (Tukey p < 0.001). However, total carbohydrate concentrations were significantly affected by experiment duration (p < 0.001) and temperature (p < 0.001). Similar to total protein, *P. strigosa* total carbohydrate was reduced under elevated temperatures compared to control conditions (Tukey p < 0.001). Additionally, the interaction between experimental duration and temperature had a significant effect on *P. strigosa* carbohydrates (p = 0.002), with *P. strigosa* exhibiting increasing carbohydrate concentrations between T_30_ and T_95_ under control temperatures (Tukey p = 0.004). In contrast, under elevated temperatures, *P. strigosa* exhibited no change in carbohydrate concentrations over time (all Tukey p > 0.05), although a trend of decreasing carbohydrate concentrations was observed from T_0_ to T_95_ (Figure 3D).

### (3.3) Effects of thermal and acidification stress on symbiont physiology of *S. siderea* and *P. strigosa*

#### (3.3.1) Symbiont physiology of *S. siderea* (Chl *a* concentration, symbiont cell density)

Experiment duration (p < 0.001), temperature (p < 0.01), and *p*CO_2_ (p = 0.04) had significant effects on *S. siderea* symbiont cell density (Figure 4A), with cell densities decreasing from T_0_ to T_95_ (Tukey p < 0.001). Additionally, *S. siderea* cell densities were reduced under elevated temperatures compared to control conditions (Tukey p = 0.017) and under extreme *p*CO_2_ compared to present day conditions (Tukey p = 0.043). The interaction between experiment duration and *p*CO_2_ also affected *S. siderea* symbiont cell density (p = 0.005); however, within *p*CO_2_ treatments the only significant change in cell densities was a decrease at low *p*CO_2_ between T_30_ and T_95_ (Tukey p = 0.003). Both experiment duration (p < 0.001) and *p*CO_2_ (p < 0.001) had significant effects on *S. siderea* Chl *a* concentration (Figure 4C). In contrast to symbiont cell density, Chl *a* concentration increased from T_0_ to T_95_ (Tukey p < 0.001). Although corals under present day and next century *p*CO_2_ treatments exhibited similar Chl *a* concentrations, corals under extreme *p*CO_2_ had significantly less Chl *a* compared to those under both present day (Tukey p = 0.002) and next century (Tukey p = 0.01) *p*CO_2_ conditions.

**Figure 4.**
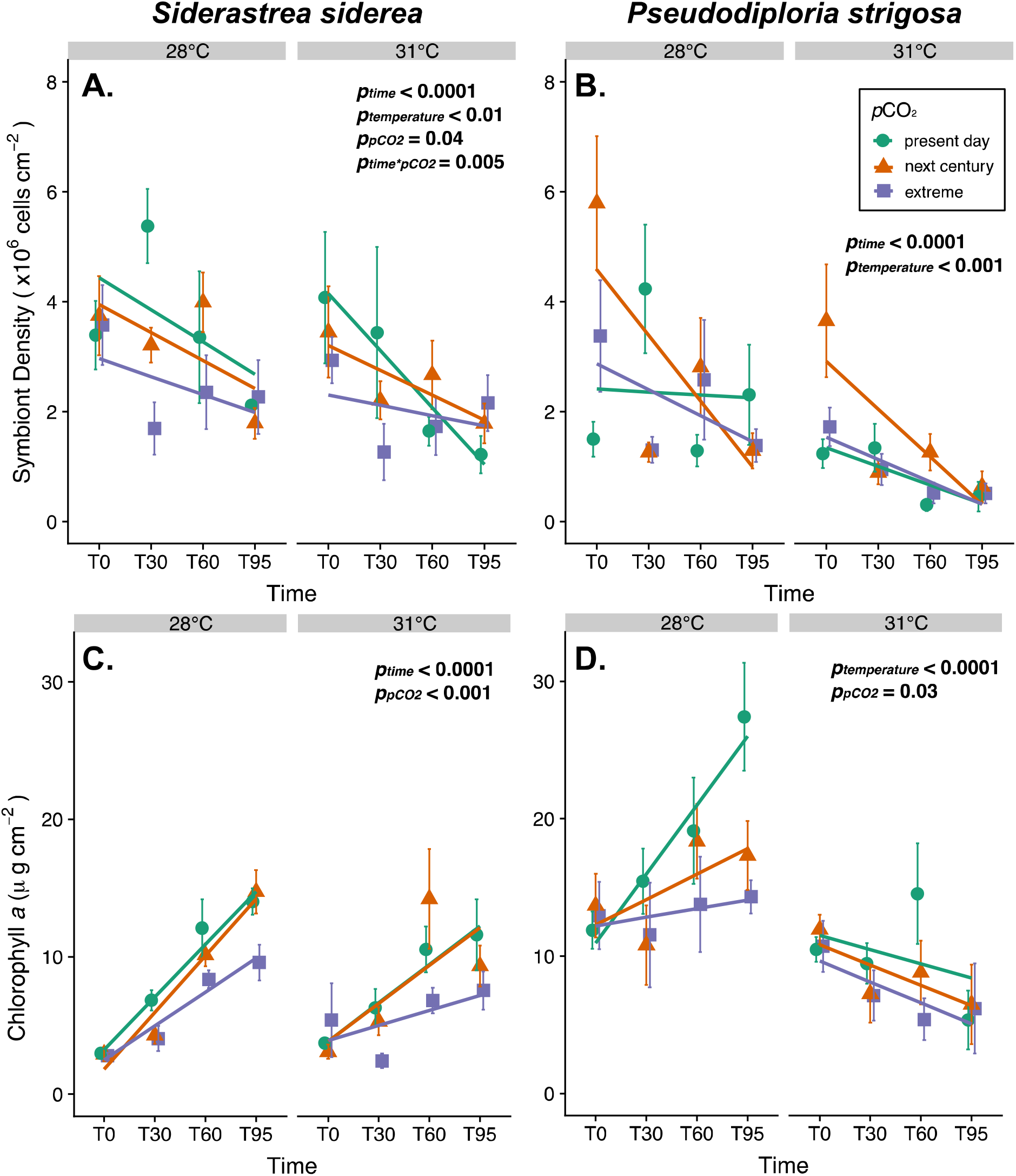
Symbiodiniaceae physiology (symbiont cell density **[A,B]** and Chl *a* concentration **[C,D]**) of *Siderastrea siderea* (**A,C**) and *Pseudodiploria strigosa* **(B,D)** across four experimental durations (day 0 = T_0_; day 30 [short-term] = T_30_; day 60 [moderate-term] = T_60_; day 95 [long-term] = T_95_). Within each panel, results are faceted by temperature treatment (28°C and 31°C) and colored by *p*CO_2_ treatment (present day [~400 μatm] = green, next century [~640 μatm] = orange, extreme [~2800 μatm] = purple). Points represent averages of n = 4-6 (*S. siderea*) and n = 3-6 (*P. strigosa*) fragments from nearshore and forereef reef zones (n = 1 / genotype). Significant factors are indicated in each panel. Lines represent linear fits (using ggplot2’s *stat_smooth()* method to visualize differences regardless of model) for each treatment through time, and error bars are standard error at each time point.

#### (3.3.2) Symbiont physiology of *P. strigosa* (Chl *a* concentration and symbiont cell density)

Experiment duration (p < 0.001) and temperature (p < 0.001) both had significant effects on *P. strigosa* symbiont cell density. Regardless of *p*CO_2_ treatment, *P. strigosa* under elevated temperatures had reduced symbiont densities compared to corals in control temperatures (Tukey p < 0.001). Additionally, *P. strigosa* symbiont density was reduced at T_95_ relative to T_0_ (Tukey p < 0.001). Temperature (p < 0.001) and *p*CO_2_ treatments (p = 0.028) had significant effects on *P. strigosa* Chl *a* concentration; however, experimental duration did not. Similar to symbiont density, *P. strigosa* exhibited reduced Chl *a* concentrations under elevated temperatures compared to control temperatures, regardless of *p*CO_2_ treatment (Tukey p < 0.001). Additionally, *P. strigosa* Chl *a* concentrations were reduced under extreme *p*CO_2_ compared to present day *p*CO_2_ treatment regardless of temperature treatment (Tukey p = 0.02).

### (3.3) Holobiont physiology through time

#### (3.3.1) *Siderastrea siderea* holobiont physiology

Overall *S. siderea* holobiont physiology clustered more strongly by *p*CO_2_ than by temperature (Figure 5A-C). There was a nearly significant effect of *p*CO_2_ on holobiont physiology after short-term exposure (T_30,_ Adonis p = 0.054; Figure 5A), and a significant effect at T_95_ (long-term: Adonis p = 0.002; Figure 5C). At T_95_, the interaction of *p*CO_2_ and temperature was also significant (Adonis p = 0.001; Figure 5C). Comparing PCAs in Figure 5A-C with individual physiology results (Figures 3A,C and 4A,C) shows that *p*CO_2_ significantly reduced *S. siderea* calcification, symbiont density, and Chl *a* concentration, but did not have a significant effect on total carbohydrates or protein. These results are consistent with the PCA loadings for calcification, symbiont density, and Chl *a* concentration discriminating between clusters of fragments in extreme *p*CO_2_ treatment and the other acidification treatments (Figure 5A-C). Reef zone did not have a significant main or interactive effect on *S. siderea* holobiont physiology for any exposure duration.

**Figure 5.**
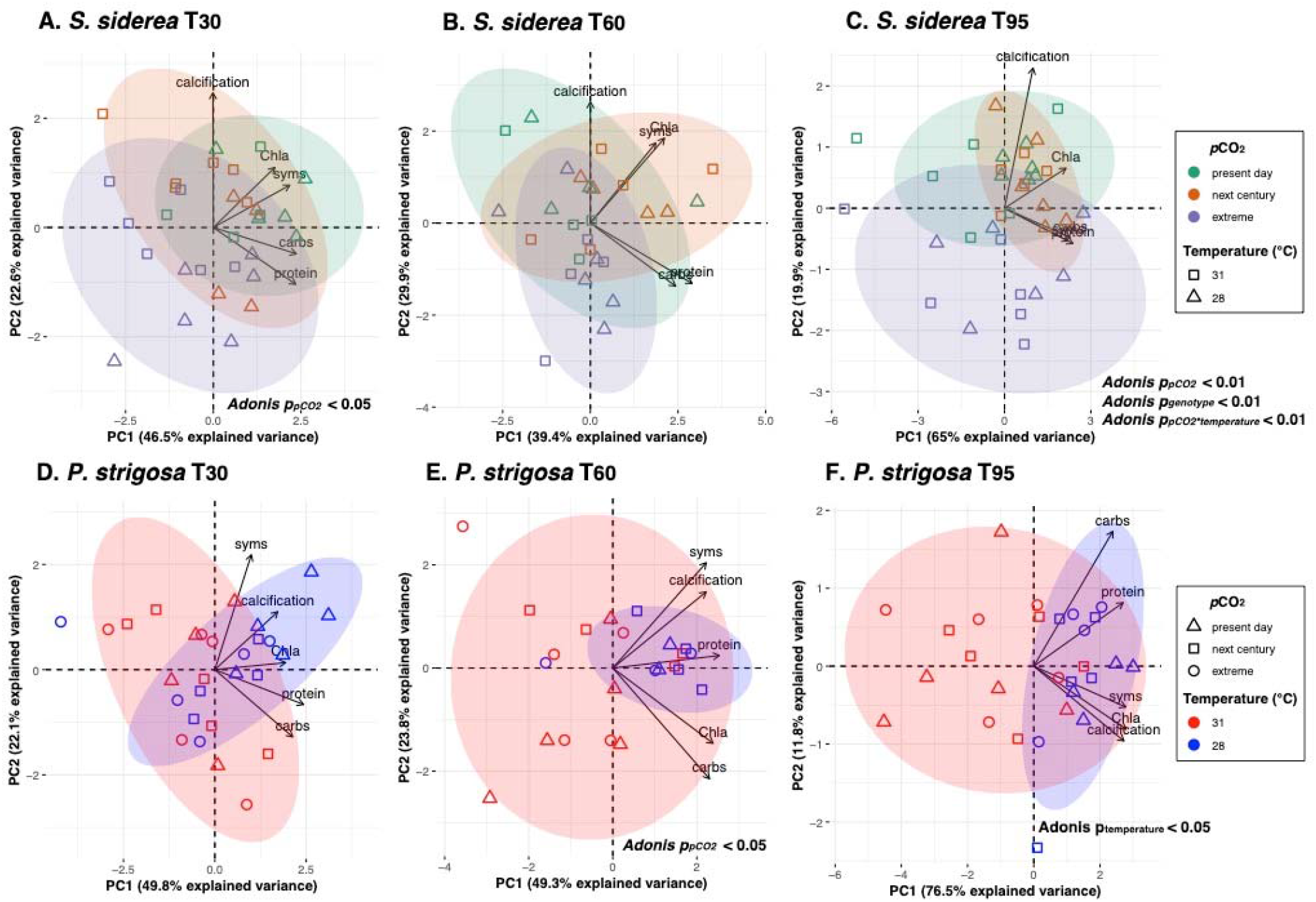
Influence of global change stressors and exposure duration on holobiont physiology. Principal Components Analysis (PCA) of log-transformed holobiont physiology data, including total carbohydrate (carbs; mg cm^−2^), total protein (protein; mg cm^−2^), symbiont density (syms; cells cm^−2^), chlorophyll *a* concentration (Chla; ug cm^−2^), and calcification rate (mg cm^−2^ day^−1^) for *Siderastrea siderea* **(A-C)** and *Pseudodiploria strigosa* **(D-F)**. Colors represent *p*CO_2_ treatment for *S. siderea* (**A-C**: green = present day [~400 μatm], orange = next century [~640 μatm], purple = extreme [~2800 μatm]) and temperature treatment for *P. strigosa* (**D-F**: red = 31°C, blue = 28°C). Shapes represent temperature treatment for *S. siderea* (**A-C**: square = 31°C, triangle = 28°C) and *p*CO_2_ treatment for *P. strigosa* (**D-F**: triangle = present day [~400 μatm], square = next century [~640 μatm], circle = extreme [~2800 μatm]). Points represent an individual coral fragment’s combined physiology at each time point (**A,D**= short-term [T_30_], **B,E**= moderate-term [T_60_], **C,F**= long-term [T_95_]). Individuals were only included if they had data for each of the five parameters at each time point. The x- and y-axes indicate the variance explained (%) by the first and second principle component, respectively.

#### (3.3.2) *Pseudodiploria strigosa* holobiont physiology

Holobiont physiology of *P. strigosa* clustered more strongly by temperature than by *p*CO_2_ treatment, especially after long-term exposure (T_95_; Figure 5D-F). At T_60_ (moderate-term), there was a significant effect of *p*CO_2_ on holobiont physiology (Adonis p = 0.029; Figure 5E). However, at T_95_ (long-term) the effect of *p*CO_2_ was no longer significant, and only temperaturehad a significant effect (Adonis p = 0.045; Figure 5F). Additionally, the interaction of reef zone and temperature had a marginally significant effect on holobiont physiology after long-term exposure (T_95_; Adonis p = 0.053; Figure 5F). Comparing PCAs in Figure 5D-F with results from individual physiology parameters (Figures 3B,D and 4B,D) shows that elevated temperature had consistent negative effects on all physiology parameters.

#### (3.3.3) Combined species holobiont physiology

*Siderastrea siderea* and *P. strigosa* had distinct holobiont physiologies at all experimental durations (Adonis p_species_ < 0.001 for short-term [T_30_], moderate-term [T_60_], and long-term [T_95_]; Figure S2; Table S5). Although species had a significant main effect on combined physiology through time, *S. siderea* and *P. strigosa* exhibit the most divergent physiologies at T_30_, and then converge to be entirely overlapping at T_60_ and T_95_ (Figure S2). There were also significant independent effects of temperature and *p*CO_2_ on the combined physiology for both species at each time point (Adonis p < 0.05 for all time points; Figure S2, Table S5).

### (3.4) *Pseudodiploria strigosa* trait correlations

*Pseudodiploria strigosa* correlation matrices (Figure S3) revealed that calcification rate was significantly correlated with all other physiology parameters (p < 0.05) after 95 days of experimental treatment (long-term; T_95_). As illustrated in *P. strigosa* results above, temperature had a main effect on the relationship between calcification and all other predictor variables (total carbohydrate and protein, symbiont density, and Chl *a* concentration; Figure 6). Correlations with calcification rate were significantly different across temperature treatment for total protein (p = 0.04) and symbiont density (p = 0.001) in *P. strigosa* at T_95_ (long-term exposure). Under elevated temperatures at T_95_, *P. strigosa* fragments with higher total protein (p = 0.04) and symbiont densities (p = 0.001) were able to maintain faster calcification rates (Figure 6 B,C). A similar trend was observed for total carbohydrates; however, this relationship was not significant (p = 0.06; Figure 6A). The interactive effect of temperature and the predictor variables on *P. strigosa* calcification rate was not significant until the final observation point of the experiment (T_95_; Figure S4). *Siderastrea siderea* correlation matrix and linear regression analyses did not reveal any significant interactions with treatment.

**Figure 6.**
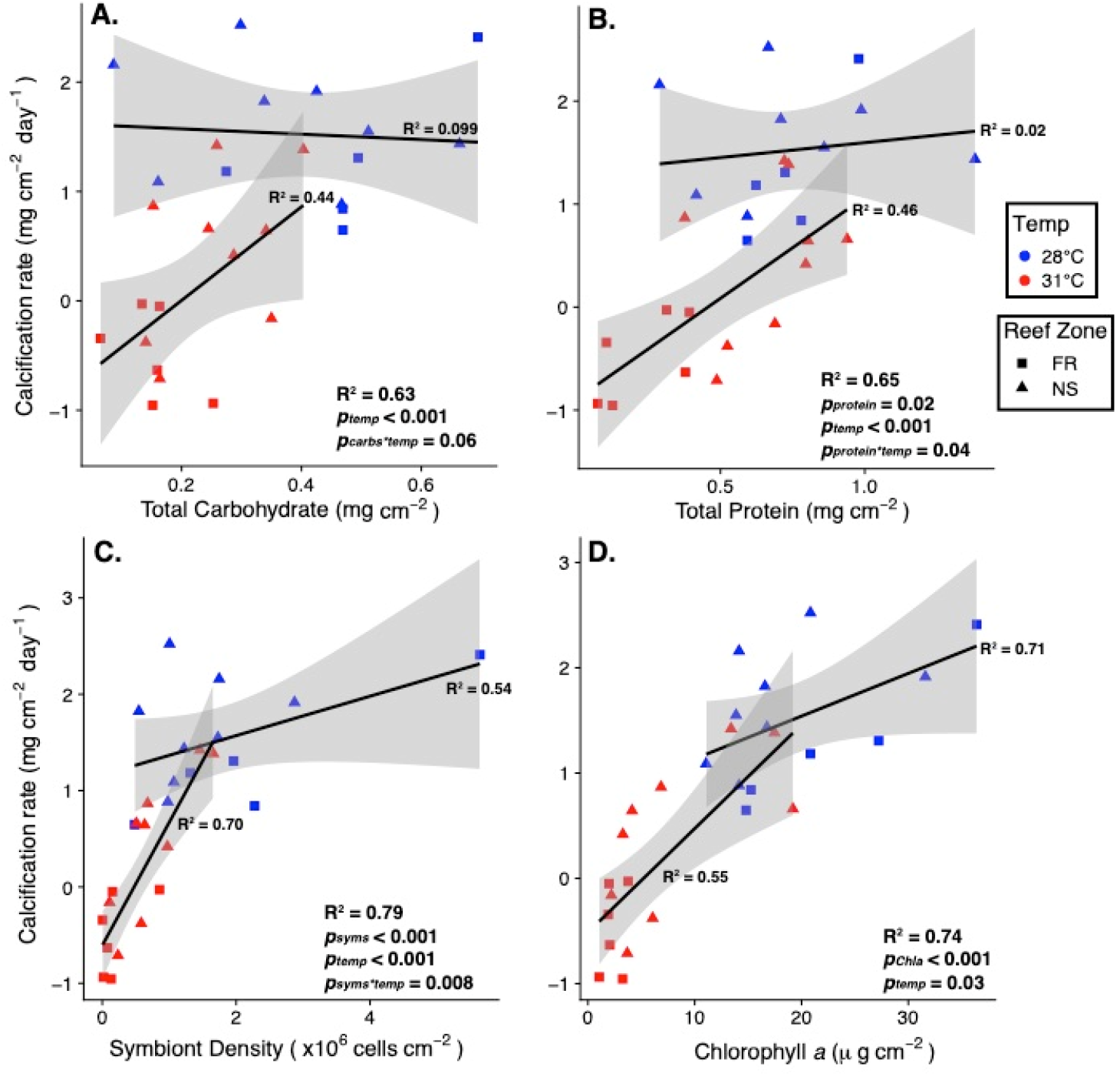
Correlations of *Pseudodiploria strigosa* calcification rate with total host carbohydrate **(A),** total host protein **(B)**, symbiont density **(C)**, and chlorophyll *a* concentration **(D)** after long-term exposure to experimental treatments (T_95_). Colors represent temperature treatment (red = 31°C, blue = 28°C) and shapes represent reef zone (square = forereef [FR], triangle = nearshore [NS]). Points represent individual coral fragments. Significant factors are indicated within each panel. Lines represent linear models of measured parameters within treatment through time, fit using ggplot2’s *stat_smooth()* method with gray shading representing 95% confidence intervals for each temperature. Conditional R-squared (R^2^) values (Nakagawa and Schielzeth, 2013) are reported for the whole model (bottom right corner of each facet) and for each temperature (next to the line of best fit).

## 4. Discussion

### (4.1) *Siderastrea siderea* and *P. strigosa* exhibit divergent responses to warming and acidification

Our results demonstrate that *S. siderea* and *P. strigosa* exhibit divergent responses to two co-occurring global change stressors—warming and acidification— and that these responses are modulated by exposure duration (i.e. short-term = T_0_-T_30_, moderate-term = T_30_-T_60_, and long-term = T_60_-T_95_). Overall physiological performance of *S. siderea* was more negatively affected by acidification through time (Figure 5A-C), while temperature stress had a more negative effect on *P. strigosa* physiology through time (Figure 5D-F). Such species-specific responses to temperature and acidification are not uncommon in reef-building corals. For example, when testing how twelve Caribbean coral species responded to crossed temperature and acidification conditions for six weeks, Okazaki et al. (2017) observed some species exhibited no growth response to either temperature or acidification (including *S. siderea* and *P. strigosa*), while other, more abundant Florida species (e.g. *O. faveolata* and *P. astreoides*) exhibited decreased calcification rates under both stressors. In contrast to Okazaki et al. (2017), we find that *S. siderea* and *P. strigosa* holobiont physiology was significantly affected by acidification and elevated temperature, respectively. Although this difference may be due to the duration of experimental stress (>30 days longer than Okazaki et al. (2017)), it could also be a result of Okazaki et al. (2017) using less extreme treatments (30.3°C and 1300 μatm *p*CO_2_) than those used here (31°C and 2800 μatm). It is also possible that populations of *S. siderea* and *P. strigosa* in the Florida Keys, the source of corals used in the Okazaki et al. (2017) study, could be less susceptible to stress than populations from the southern MBRS used here. According to the climate variability hypothesis (CVH; Stevens, 1989), higher latitude populations (e.g. Florida Keys) that experience more variable thermal regimes are predicted to be more phenotypically flexible and exhibit a wider range of thermal tolerances compared to populations existing closer to the equator (e.g. MBRS; Bozinovic et al., 2011). In a meta-analysis of Caribbean coral calcification responses to acidification, elevated temperature, and the combination of the two stressors, Bove et al. (2020) found similar regional differences in stress responses between corals from the Florida Keys and Belize. While calcification of Florida Keys corals did not clearly respond to acidification, elevated temperature, or their combination, elevated temperature reduced calcification rates in corals from Belize (Bove et al., 2020). While acknowledging differences in annual temperature variability, Bove et al. (2020) highlight differences in experimental treatment extremes as the main driver of calcification. While consideration of treatment level is critical, such population-level differences in stress tolerance have been previously observed in corals, for example in *Acropora millepora* across 5° of latitude on the Great Barrier Reef (Dixon et al., 2015). Regardless, results of the present study contribute to a growing body of literature supporting *S. siderea*’s resistance to conditions of elevated temperature and acidification (Banks and Foster, 2016; Bove et al., 2019; Castillo et al., 2014; Castillo et al., 2011; Davies et al., 2016).

High resistance of *S. siderea* to global change stressors was previously reported by Castillo et al. (2014), which found that only the most extreme temperature (32°C) and acidification (2553 μatm *p*CO_2_) treatments resulted in reduced calcification rates. In the context of global change scenarios projected by the IPCC, Castillo et al. (2014) concluded that *S. siderea* will be more negatively impacted by elevated temperatures over the coming century, given that the IPCC’s next-century acidification projections did not reduce calcification rates. The findings of the present study are consistent with this previous work, as only the extreme *p*CO_2_ treatment– but not next century *p*CO_2_ treatment–reduced calcification rate in *S. siderea* (Figure 2A). Gene expression profiling of *S. siderea* from the Castillo et al. (2014) coral fragments revealed that thermal stress caused large-scale down regulation of gene expression, while acidification stress elicited upregulation of proton transport genes (Davies et al., 2016), potentially offsetting the effects of acidification at the site of calcification (e.g. Ries, 2011; Schoepf et al., 2017). These findings provide further support for *S. siderea*’s ability to acclimate to acidification stress.

Bove et al. (2019) investigated the combined effects of similar temperature and acidification treatments on four species of reef building corals: *S. siderea, P. strigosa, Porites astreoides,* and *Undaria tenuifolia.* After 93 days, all species exhibited calcification declines under increased *p*CO_2_. However, *P. strigosa* was the only species that exhibited reduced calcification under elevated temperature, which is consistent with results present here and highlights that thermal stress more negatively impacts *P. strigosa* than *S. siderea* (Figure 5D-F). Bove et al. (2019) also found that *S. siderea* was the most resistant of the four species studied, as it was able to maintain positive calcification rates even in the most extreme acidification treatment (~3300 μatm *p*CO_2_)—findings that are also corroborated here (Figure 2A).

Interestingly, by quantifying net calcification rates at 30-day increments, we show that *S. siderea* net calcification was negative under extreme *p*CO_2_ at T_60_ and then these rates recovered by T_95_ (Figure 2A). This suggests that these corals are acclimating to stressful conditions over time, perhaps through transcriptome plasticity, as previously proposed by Davies et al. (2016).

### (4.2.) Stress differentially modulates coral physiology across species

Under thermal and acidification stress, corals can draw on energy reserves, including lipids, proteins, and carbohydrates, to maintain and/or produce tissue and skeleton (Anthony et al., 2009; Schoepf et al., 2013). In addition to using energetic reserves, heterotrophic feeding (Aichelman et al., 2016; Drenkard et al., 2013; Edmunds, 2011; Towle et al., 2015) or enhanced productivity of Symbiodiniaceae owing to CO_2_ fertilization of symbiont photosynthesis (Brading et al., 2011) can augment energetic resources in zooxanthellate corals. Coral energetic reserves can therefore influence resistance to and recovery from stress events (Edmunds et al., 2016; Grottoli et al., 2006; Grottoli et al., 2014; Schoepf et al., 2015).

Results of the present study show that the host energy reserves (protein, carbohydrates) of *S. siderea* and *P. strigosa* responded to global change stress (warming, acidification) in different ways. Between T_0_ and T_60_, *P. strigosa* exhibited reduced carbohydrates regardless of treatment, indicating catabolism of this energy reserve (Figure 3D). This was followed by the restoration of carbohydrates (i.e. acclimation) at control temperatures at T_95_ (Figure 3D), which likely supported the positive calcification rates also observed under these conditions (Figure 6A, Figure S4). Protein reserves do not show the same trend as carbohydrates through time (Figure 3B), potentially owing to *P. strigosa* catabolizing carbohydrates before proteins, which has been observed over shorter time scales for other scleractinian coral species (Grottoli et al., 2004). However, elevated protein reserves did predict faster calcification rates in *P. strigosa* under elevated temperatures, but not until the final observation point of the experiment (T_95_; Figure 6B, Figure S4). As photosynthate translocated from symbionts is a major source of carbohydrates to the coral host (Burriesci et al., 2012; Muscatine, 1990), reductions in symbiont cell density, Chl *a* concentration, and carbohydrates of *P. strigosa* at elevated temperature suggest that symbionts were translocating fewer resources to the host, which likely contributed to observed reductions in calcification under these elevated temperature conditions, particularly after long-term exposure (T_95_, Figure 6). Total protein and carbohydrate abundances of *S. siderea*, similar to those of *P. strigosa,* declined under elevated temperatures (Figure 3A,C)—consistent with previous work highlighting upregulation of protein catabolism pathways in *S. siderea* exposed to long-term thermal stress (Davies et al., 2016).

An overall trend in reduced symbiont density and increased Chl *a* concentration through time were observed under most *p*CO_2_ and temperature conditions, except for *P. strigosa* under elevated temperature (Figure 4). Given that both species under most treatments exhibited this pattern, it cannot be ruled out that these changes in symbiont physiology were influenced by other factors, including incomplete symbiont acclimation to experimental light environment (Roth, 2014) and seasonal patterns in symbiont density and pigment concentration (Fitt et al., 2000)—which may have masked the symbiont response to thermal stress within *S. siderea*. In contrast, *P. strigosa* exhibited reduced symbiont density and Chl *a* concentrations under elevated temperature (Figure 4B,D), a pattern that is more consistent with thermally induced bleaching (Brown, 1997; Fitt et al., 2001; Glynn, 1993; Warner et al., 1999; Weis, 2008) and further illustrates the susceptibility of this species to thermal stress.

### (4.3.) Nearshore *P. strigosa* are more resistant than forereef conspecifics

Natal reef zone was a significant predictor of host physiology, particularly for *P. strigosa*, as nearshore corals calcified faster overall (Figure 2B). Although reef zone differences in calcification were observed for *S. siderea* (particularly through time), colonies from one reef zone did not clearly outperform colonies from the other reef zone (Figure 2A). In contrast, the reef-zone-specific calcification response of *P. strigosa* may arise from local adaptation of the host to distinct temperature regimes. On the MBRS, nearshore habitats are characterized by higher maximum temperatures, greater annual temperature range, and more days above the regional thermal bleaching threshold compared to forereef sites (Baumann et al., 2016). Local adaptation to distinct reef zones is not uncommon in corals, and has been previously shown to affect responses of corals to thermal stress. For example, *P. astreoides* was previously found to be locally adapted to distinct thermal regimes in the Florida Keys, with inshore corals demonstrating higher thermal tolerance, constitutively higher expression of specific metabolic genes, and greater gene expression plasticity compared to offshore conspecifics (Kenkel et al., 2013a; Kenkel et al., 2013b; Kenkel and Matz, 2016). Additionally, *P. strigosa* is a hermaphroditic broadcast spawning species, and previous work on populations from the Flower Garden Banks demonstrated that larvae have short pelagic larval durations (PLD; Davies et al., 2017), which could facilitate local adaptation as larvae are more likely to locally recruit (Davies et al., 2017; Mayorga-Adame et al., 2017). However, it is unknown if *P. strigosa* on the MBRS have similarly short PLDs. We hypothesize that nearshore *P. strigosa* are locally adapted and/or acclimated to more variable and stressful nearshore conditions, allowing maintenance of higher calcification rates under thermal stress compared to their forereef counterparts.

It is important to note that responses to stress based on reef zone could be obscured by uneven sampling across reef zones, as forereef genotypes of *P. strigosa* were slightly underrepresented in the experiment (2 genotypes present vs. the standard 3) due to mortality before the experiment began. It is possible that full replication of genotype would have yielded different effects of reef zone in other *P. strigosa* physiology parameters.

Although the *S. siderea* PLD is unknown, previous work has shown high population connectivity across 2,000 km of the Brazil coast (Nunes et al., 2011). If MBRS *S. siderea* populations are similarly connected, it is possible that no genetic differences exist across nearshore and forereef environments—potentially explaining the lack of reef-zone-specific responses observed here. Yet this result is inconsistent with previous findings that forereef colonies of MBRS *S. siderea* exhibited greater physiological stress than inshore colonies when exposed to higher temperatures (Castillo and Helmuth, 2005), reduced skeletal extension rates relative to inshore colonies over multi-decadal warming of the reef system (Castillo et al., 2012), and reduced symbiont photophysiology relative to inshore colonies under higher temperatures (Davies et al., 2018). However, consistent with results presented here, Bove et al. (2019) also found no evidence for reef-zone differences in physiology for *S. siderea,* or for any of the other species tested. Nevertheless, the observation that both nearshore and forereef *S. siderea* performed well under global change stressors provides further support for resistance of this species.

### (4.4.) Time-course experiments reveal acclimation to thermal stress in two common Caribbean corals

This study contributes to a growing body of literature demonstrating the value of assessing time-course physiology of corals exposed to global change stressors. Although studies investigating independent effects of temperature and acidification on scleractinian corals have yielded great insight into the effects of future global change on coral systems (e.g. Albright et al., 2018; Anthony et al., 2011; Carricart-Ganivet et al., 2012; Comeau et al., 2013; Jokiel and Coles, 1990; Jury et al., 2010), the combined effects of these stressors remain less explored— particularly in the context of how the coral stress response is modulated by the duration of those stressors. By characterizing coral host and symbiont physiology of a colony through time, acclimatory responses were identified in two common Caribbean reef-building coral species. The results of this study provide further evidence of the species-specific nature of this acclimation.

For example, under extreme *p*CO_2_ and elevated temperature, *S. siderea* calcification rate appears to recover by the end of the experiment while *P. strigosa* calcification rate declines into negative net calcification. In addition to furthering our understanding of how corals could respond to projected future ocean conditions, the exposure duration component of this study suggests that species will exhibit differential persistence through ephemeral stress events. It is clear that local heat waves that raise SST and upwelling events that reduce pH, which already threaten coral populations, may threaten coral species in different ways in the future depending on the timescales of these events.

Acclimation is an important mechanism by which corals can withstand changing environmental conditions, and transcriptome plasticity is one way by which corals can acclimate to stress (Davies et al., 2016; Kenkel and Matz, 2016). A coral reciprocal transplant study between reef zones in the Florida Keys demonstrated that adaptive gene expression plasticity, specifically plasticity of stress response genes, was associated with reduced susceptibility to a summer bleaching event (Kenkel and Matz, 2016). In addition to plasticity providing a mechanism for acclimation within a generation, rapid evolutionary adaptation of corals to warmer oceans has also been observed on the Great Barrier Reef (Dixon et al., 2015; Matz et al., 2018). However, recent widespread declines in coral abundance, diversity, and health suggest that rates of intra- and trans-generational adaptation to global change stressors within most coral populations are insufficient for mitigating the deleterious impacts of recent and future CO_2_-induced global change (Thomas et al., 2018). Understanding the interplay of acclimation and adaptation in scleractinian corals is therefore essential for projecting how coral reef ecosystems will fare in the higher-CO_2_ future (Chevin et al., 2010; Thomas et al., 2018). This study furthers understanding of how exposure duration modulates coral physiology across reef zones in two prevalent Caribbean reef-building species. Future studies focusing on long-term acclimation capacities of corals will further elucidate mechanisms of resistance and resilience in corals’ response to global stressors.

## 5. Acknowledgements

We thank the Belize Fisheries Department for all coral collection permits. Isaac Westfield, Amanda Dwyer, Sara Williams, and Louise Cameron are acknowledged for assisting with experimental maintenance at Northeastern University. Thanks to both the Marchetti and Septer labs at the University of North Carolina at Chapel Hill (UNC) for assistance with equipment and lab space to complete the physiology assays. We also thank Samir Patel, Savannah Swinea, Bailey Thomasson, Forrest Buckthal, and Cori Lopazanski for assisting with preparing corals for physiology assays at UNC.

## 6. Funding

This project was supported by NSF award OCE-1437371 (to J.B.R.), NSF award OCE-1459522 (to K.D.C.), NOAA award NA13OAR4310186 (to J.B.R. and K.D.C.), and the National Academies of Sciences, Engineering, and Medicine Gulf Research Program Fellowship (to S.W.D.). H.E.A. was supported by an NSF GRFP (2016222953) and S.W.D. was a Simons Foundation Fellow of the Life Sciences Research Foundation (LSRF) while completing part of this research.

## Notes

https://github.com/hannahaichelman/TimeCourse_Physiology

### Competing Interest Statement

The authors have declared no competing interest.

