## Supplemental Information for "Exposure duration modulates the response of Caribbean corals to global change stressors"

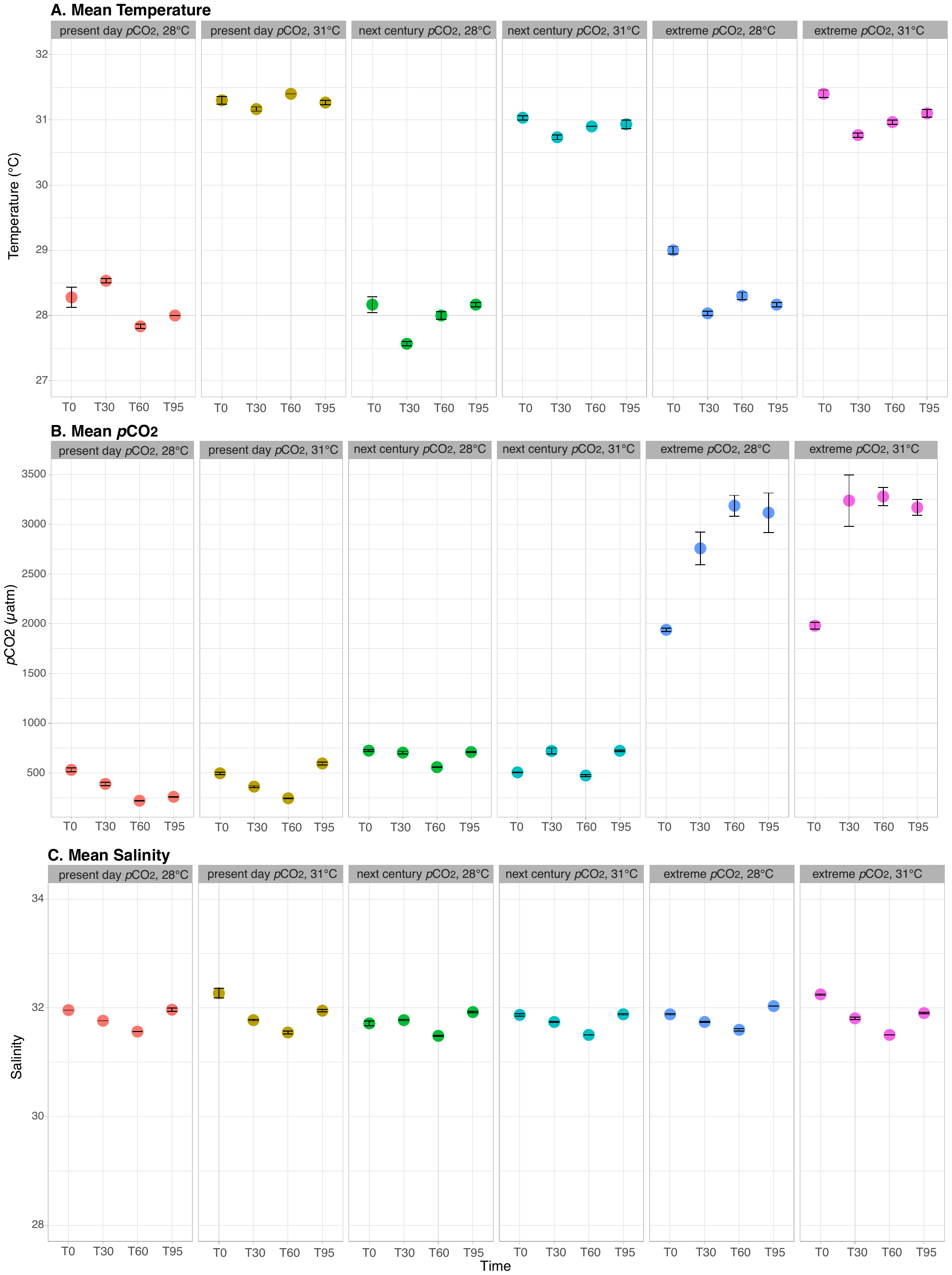

**Figure S1.** Water quality parameters measured at each time point (short-term = T_30_; moderate-term = T_60_; long-term = T_95_) of the experiment, including temperature **(A)**, partial pressure of carbon dioxide (*p*CO_2_, **B**), and salinity **(C)**. Facets represent each of the six treatments (*p*CO_2_: present day [~400 μatm], next century [~640 μatm], extreme [~2800 μatm]; temperature: 28°C, 31°C). Points are mean $\pm$ standard error of three replicate measurements.

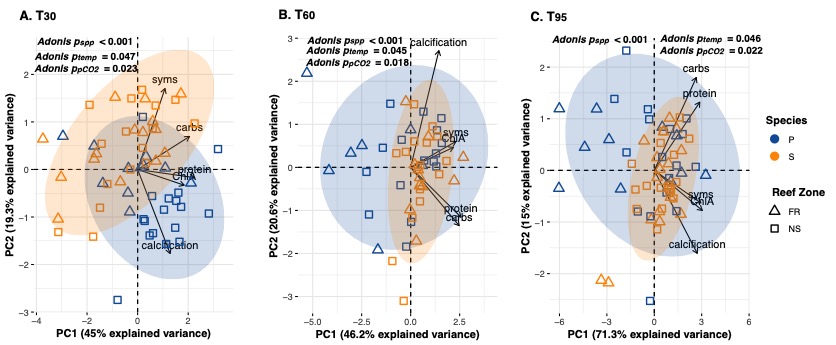

**Figure S2.** Principal Components Analysis (PCA) of *Siderastrea siderea* and *Pseudodiploria strigosa* log-transformed physiology data, including total carbohydrate (carbs; mg cm^-2^), total protein (protein; mg cm^-2^), Symbiodiniaceae density (syms; cells cm^-2^), chlorophyll *a* concentration (ChlA; μg cm^-2^), and calcification rate (mg cm^-2^ day^-1^). Colors represent species (*S. siderea* = orange, *P. strigosa* = blue) and shapes represent reef zone (square = forereef [FR], triangle = nearshore [NS]). Points represent an individual coral fragment’s combined physiology after each experimental duration (**A** = short-term [T_30_], **B** = moderate-term [T_60_], **C** = long-term [T_95_]). Individuals were only included if they had a measure for each of the five parameters at that time point. The x- and y-axes indicate the variance explained (%) by first and second principle components, respectively.

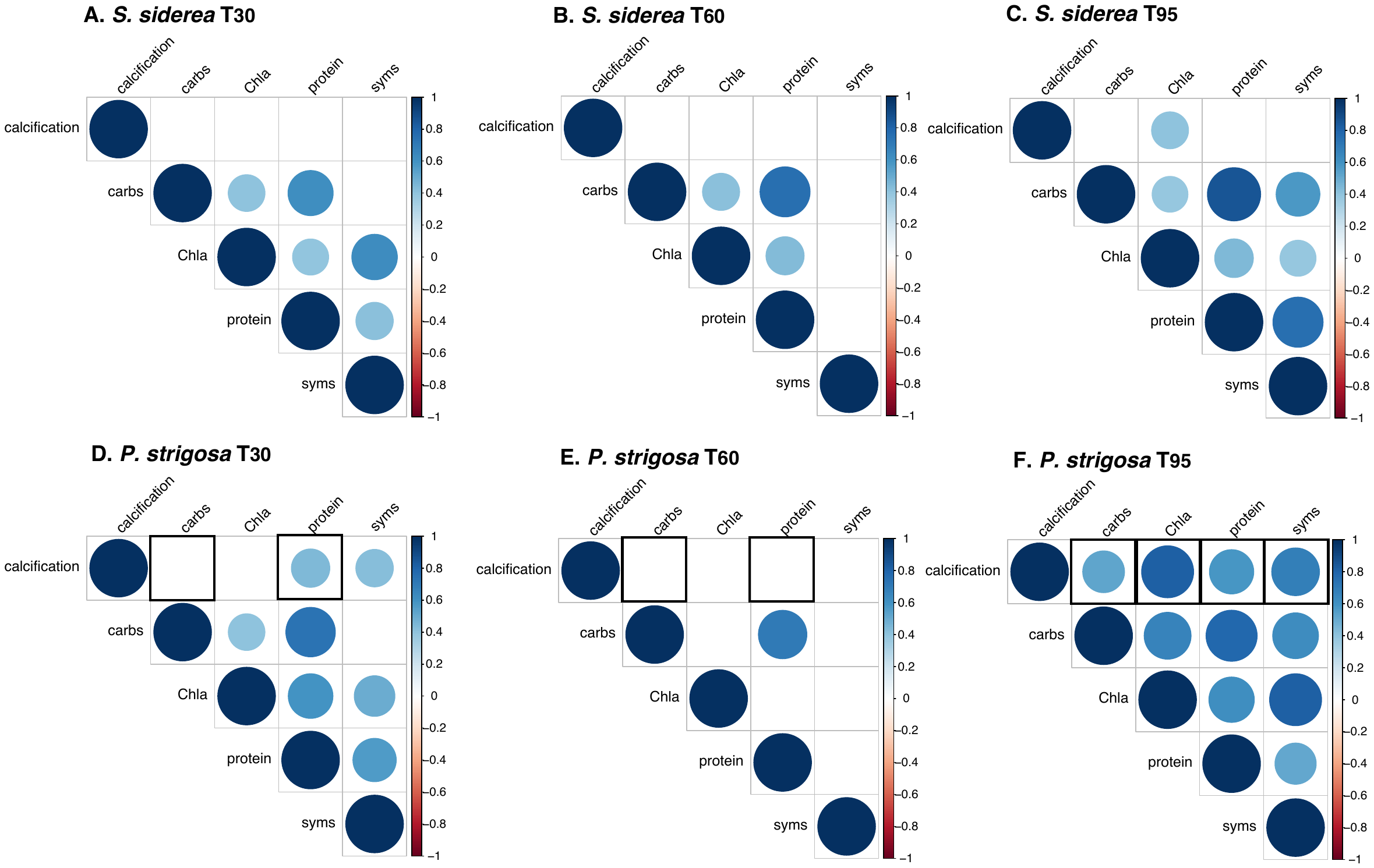

**Figure S3.** Correlation matrices for *S. siderea* (**A-C**) and *P. strigosa* (**D-F**) host and symbiont physiology parameters, including total carbohydrate (carbs; mg cm^-2^), total protein (protein; mg cm^-2^), Symbiodiniaceae density (syms; cells cm^-2^), chlorophyll *a* concentration (Chla; μg cm^-2^), and calcification rate (mg cm^-2^ day^-1^) through time (**A,D** = short-term [T_30_], **B,E** = moderate-term [T_60_], **C,F** = long-term [T_95_]). Positive correlations are represented by blue colors and negative correlations are represented by red colors. Circle color intensity and size are proportional to the correlation coefficients. Insignificant correlations (p > 0.05) are blank. Black boxes indicate correlations that are specifically discussed in the main manuscript (Figure 6) and supplementary information (Figure S4).

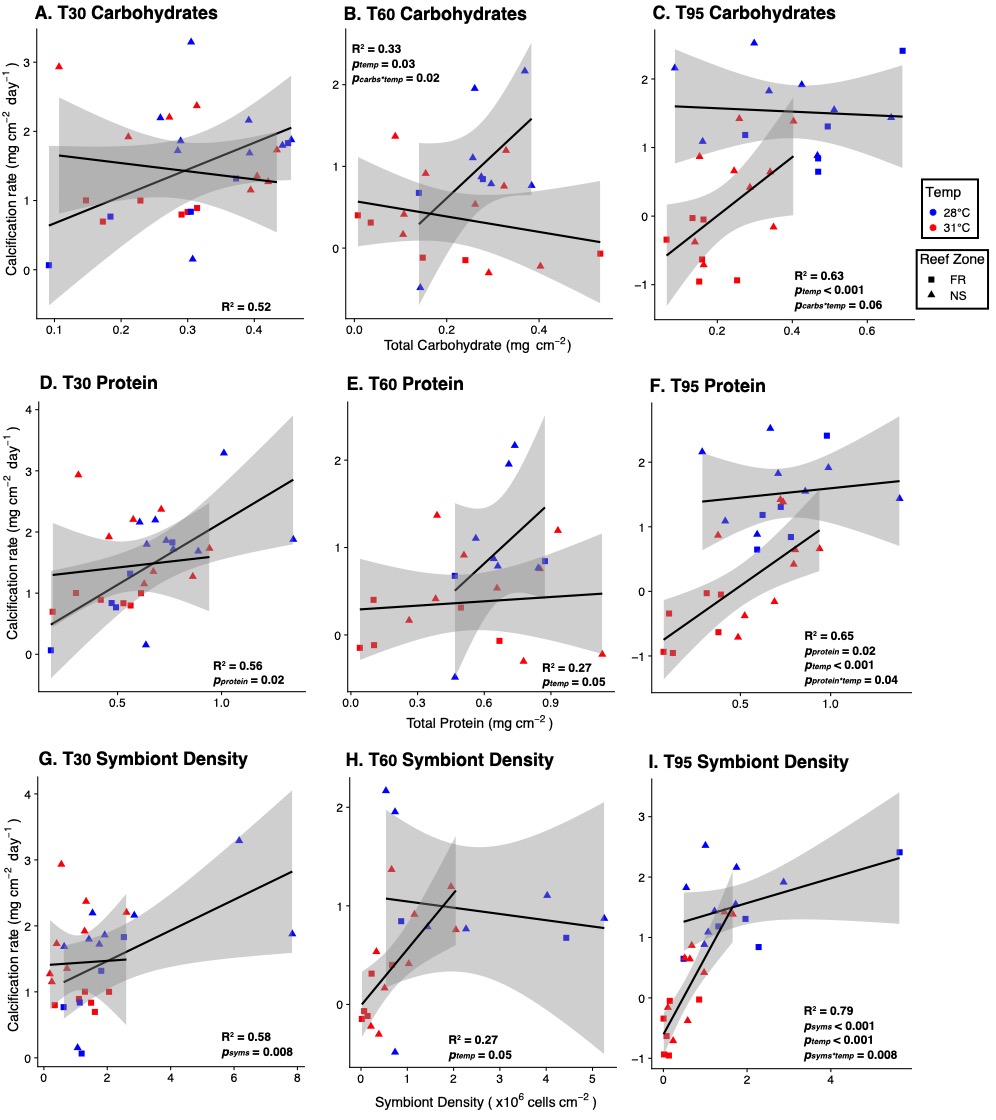

**Figure S4.** Correlations of *Pseudodiploria strigosa* calcification with total carbohydrate (**A-C**), total protein (**D-F**), and symbiont density (**G-I**). Points represent an individual coral fragment at each time point (**A,D,G** = short-term [T_30_], **B,E,H** = moderate-term [T_60_], **C,F,I** = long-term [T_95_]). Colors represent temperature treatment (red = 31°C, blue = 28°C) and shapes represent reef zone (square = forereef [FR], triangle = nearshore [NS]). Significant factors for each panel are indicated within that panel. Lines represent linear fits between the physiology parameters by temperature (using ggplot2’s *stat_smooth()* method) with gray shading representing 95% confidence intervals for each temperature. Conditional R-squared values (R^2^) are reported.

**Table S1. Measured water quality parameters.** Average measured parameters for all experimental treatments: salinity (Sal), temperature (Temp), pH, total alkalinity (TA), and dissolved inorganic carbon (DIC). All values are displayed as average $\pm$ standard error, and n = 3 for all measurements except for Present Day *p*CO_2_, 28°C treatment at T_95_ (n = 2).

| **Treatment** | **Duration** | **Temp (°C)** | **Sal (psu)** | **TA (**$\boldsymbol{\mu}$**M)** | **pH_M-NBS_** | **DIC (**$\boldsymbol{\mu}$**M)** |
| --- | --- | --- | --- | --- | --- | --- |
| Present Day *p*CO_2_, 28°C | *T_0_* | 28.3$\pm$0.2 | 31.96$\pm$0.0 | 1889$\pm$0.0 | 8.06$\pm$0.002 | 1688$\pm$0.8 |
|  | *T_30_* | 28.5$\pm$0.0 | 31.76$\pm$0.0 | 1972$\pm$3.5 | 8.33$\pm$0.003 | 1695$\pm$7.3 |
|  | *T_60_* | 27.8$\pm$0.0 | 31.56$\pm$0.0 | 2031$\pm$0.3 | 8.19$\pm$0.002 | 1633$\pm$1.8 |
|  | *T_95_* | 28.0$\pm$0.0 | 31.96$\pm$0.033 | 2079$\pm$2.0 | 8.40$\pm$0.003 | 1700$\pm$3.9 |
| Present Day *p*CO_2_, 31°C | *T_0_* | 31.3$\pm$0.1 | 32.27$\pm$0.089 | 1880$\pm$0.0 | 8.14$\pm$0.008 | 1642$\pm$3.5 |
|  | *T_30_* | 31.2$\pm$0.0 | 31.77$\pm$0.011 | 2018$\pm$4.9 | 8.34$\pm$0.002 | 1698$\pm$5.0 |
|  | *T_60_* | 31.4$\pm$0.0 | 31.55$\pm$0.029 | 2041$\pm$0.6 | 8.18$\pm$0.009 | 1632$\pm$3.6 |
|  | *T_95_* | 31.3$\pm$0.0 | 31.95$\pm$0.022 | 2108$\pm$4.4 | 8.08$\pm$0.003 | 1863$\pm$3.9 |
| Next century *p*CO_2_, 28°C | *T_0_* | 28.2$\pm$0.1 | 31.71$\pm$0.044 | 1874$\pm$0.0 | 7.96$\pm$0.003 | 1720$\pm$1.3 |
|  | *T_30_* | 27.6$\pm$0.0 | 31.77$\pm$0.011 | 2022$\pm$5.2 | 8.03$\pm$0.005 | 1847$\pm$7.5 |
|  | *T_60_* | 28.0$\pm$0.1 | 31.48$\pm$0.011 | 2053$\pm$1.8 | 7.96$\pm$0.005 | 1832$\pm$1.9 |
|  | *T_95_* | 28.2$\pm$0.0 | 31.92$\pm$0.011 | 2121$\pm$4.3 | 8.01$\pm$0.004 | 1928$\pm$4.3 |
| Next century *p*CO_2_, 31°C | *T_0_* | 31.0$\pm$0.0 | 31.87$\pm$0.022 | 1936$\pm$0.0 | 8.14$\pm$0.003 | 1695$\pm$1.5 |
|  | *T_30_* | 30.7$\pm$0.0 | 31.74$\pm$0.011 | 2029$\pm$7.0 | 8.02$\pm$0.002 | 1836$\pm$8.6 |
|  | *T_60_* | 30.9$\pm$0.0 | 31.50$\pm$0.0 | 2032$\pm$1.3 | 7.98$\pm$0.007 | 1763$\pm$3.5 |
|  | *T_95_* | 30.9$\pm$0.1 | 31.88$\pm$0.011 | 2101$\pm$3.9 | 7.99$\pm$0.000 | 1896$\pm$2.6 |
| Extreme *p*CO_2_, 28°C | *T_0_* | 29.0$\pm$0.1 | 31.88$\pm$0.011 | 1952$\pm$0.0 | 7.59$\pm$0.004 | 1915$\pm$1.1 |
|  | *T_30_* | 28.0$\pm$0.0 | 31.74$\pm$0.011 | 2089$\pm$6.2 | 7.35$\pm$0.006 | 2092$\pm$5.7 |
|  | *T_60_* | 28.3$\pm$0.1 | 31.59$\pm$0.019 | 2101$\pm$2.3 | 7.29$\pm$0.009 | 2123$\pm$2.5 |
|  | *T_95_* | 28.2$\pm$0.0 | 32.03$\pm$0.0 | 2132$\pm$1.9 | 7.35$\pm$0.008 | 2148$\pm$6.9 |
| Extreme *p*CO_2_, 31°C | *T_0_* | 31.4$\pm$0.1 | 32.25$\pm$0.011 | 1751$\pm$0.0 | 7.57$\pm$0.007 | 1719$\pm$1.9 |
|  | *T_30_* | 30.8$\pm$0.0 | 31.81$\pm$0.022 | 2079$\pm$4.3 | 7.27$\pm$0.003 | 2092$\pm$6.8 |
|  | *T_60_* | 31.0$\pm$0.0 | 31.50$\pm$0.0 | 2091$\pm$1.2 | 7.23$\pm$0.005 | 2105$\pm$4.7 |
|  | *T_95_* | 31.1$\pm$0.1 | 31.90$\pm$0.011 | 2126$\pm$4.6 | 7.23$\pm$0.000 | 2132$\pm$6.5 |

**Table S2. Calculated water quality parameters.** Average calculated parameters for all treatments: *p*CO_2_ of the mixed gases in equilibrium with seawaters (*p*CO_2(gas-e)_); calculated pH (pHc); carbonate ion concentration ([CO_3_^2‒^]); bicarbonate ion concentration ([HCO_3_^‒^]); dissolved carbon dioxide ([CO_2_]_(SW)_), and aragonite saturation state ($\Omega$_A_). All values are displayed as average $\pm$ standard error, and n = 3 for all measurements except for Present Day *p*CO_2_, 28°C treatment at T_95_ (n = 2).

| **Treatment** | **Duration** | **pCO_2(gas-e)_ (**$\boldsymbol{\mu}$**atm-v)** | **pH_c-NBS_** | **[CO_3_^2-^] (**$\boldsymbol{\mu}$**M)** | **[HCO_3_^-^] (**$\boldsymbol{\mu}$**M)** | **[CO_2_]_(SW)_ (**$\boldsymbol{\mu}$**M)** | $\boldsymbol{\Omega}$**_A_** |
| --- | --- | --- | --- | --- | --- | --- | --- |
| Present Day *p*CO_2_, 28°C | *T_0_* | 531$\pm$19 | 8.03$\pm$0.01 | 143$\pm$0.5 | 1531$\pm$1.1 | 14$\pm$0.2 | 2.3$\pm$0.01 |
|  | *T_30_* | 389$\pm$17 | 8.16$\pm$0.02 | 194$\pm$5 | 1491$\pm$11.6 | 10$\pm$0.4 | 3.2$\pm$0.08 |
|  | *T_60_* | 220$\pm$2 | 8.36$\pm$0.002 | 274$\pm$1.1 | 1354$\pm$2.9 | 6$\pm$0.1 | 4.5$\pm$0.02 |
|  | *T_95_* | 260$\pm$3 | 8.32$\pm$0.003 | 264$\pm$1.1 | 1430$\pm$4.9 | 7$\pm$0.1 |  |
| Present Day *p*CO_2_, 31°C | *T_0_* | 496$\pm$12 | 8.06$\pm$0.008 | 167$\pm$2.1 | 1463$\pm$5.3 | 12$\pm$0.3 | 2.8$\pm$0.04 |
|  | *T_30_* | 362$\pm$8 | 8.20$\pm$0.008 | 225$\pm$3.2 | 1464$\pm$6.9 | 9$\pm$0.2 | 3.8$\pm$0.05 |
|  | *T_60_* | 245$\pm$5 | 8.33$\pm$0.006 | 283$\pm$2.7 | 1344$\pm$6.2 | 6$\pm$0.1 | 4.8$\pm$0.05 |
|  | *T_95_* | 595$\pm$15 | 8.04$\pm$0.01 | 180$\pm$3.5 | 1669$\pm$6.3 | 14$\pm$0.3 | 3.0$\pm$0.06 |
| Next century *p*CO_2_, 28°C | *T_0_* | 724$\pm$9 | 7.92$\pm$0.005 | 116$\pm$0.7 | 1585$\pm$1.8 | 19$\pm$0.2 | 1.9$\pm$0.01 |
|  | *T_30_* | 704$\pm$14 | 7.96$\pm$0.01 | 133$\pm$1.4 | 1695$\pm$8.5 | 18$\pm$0.4 | 2.2$\pm$0.02 |
|  | *T_60_* | 558$\pm$2 | 8.05$\pm$0.001 | 163$\pm$0.6 | 1655$\pm$2.1 | 14$\pm$0.1 | 2.7$\pm$0.01 |
|  | *T_95_* | 710$\pm$6 | 7.97$\pm$0.003 | 147$\pm$0.7 | 1762$\pm$4.2 | 18$\pm$0.1 | 2.4$\pm$0.01 |
| Next century *p*CO_2_, 31°C | *T_0_* | 506$\pm$5 | 8.06$\pm$0.003 | 172$\pm$1.0 | 1511$\pm$2.3 | 12$\pm$0.1 | 2.9$\pm$0.02 |
|  | *T_30_* | 721$\pm$30 | 7.95$\pm$0.02 | 145$\pm$4.5 | 1673$\pm$11.2 | 17$\pm$0.8 | 2.4$\pm$0.08 |
|  | *T_60_* | 474$\pm$8 | 8.11$\pm$0.01 | 193$\pm$2.0 | 1559$\pm$5.2 | 11$\pm$0.2 | 3.2$\pm$0.03 |
|  | *T_95_* | 722$\pm$7 | 7.97$\pm$0.004 | 156$\pm$1.2 | 1723$\pm$2.1 | 17$\pm$0.1 | 2.6$\pm$0.02 |
| Extreme *p*CO_2_, 28°C | *T_0_* | 1939$\pm$18 | 7.55$\pm$0.004 | 59$\pm$0.4 | 1807$\pm$1.1 | 49$\pm$0.4 | 1.0$\pm$0.01 |
|  | *T_30_* | 2758$\pm$165 | 7.43$\pm$0.03 | 47$\pm$2.9 | 1974$\pm$4.7 | 71$\pm$4.3 | 0.8$\pm$0.05 |
|  | *T_60_* | 3186$\pm$105 | 7.38$\pm$0.01 | 42$\pm$1.3 | 1999$\pm$1.3 | 81$\pm$2.6 | 0.7$\pm$0.02 |
|  | *T_95_* | 3117$\pm$198 | 7.39$\pm$0.03 | 45$\pm$2.6 | 2023$\pm$4.5 | 80$\pm$5.0 | 0.7$\pm$0.04 |
| Extreme *p*CO_2_, 31°C | *T_0_* | 1980$\pm$34 | 7.50$\pm$0.01 | 52$\pm$0.7 | 1621$\pm$1.8 | 47$\pm$0.8 | 0.9$\pm$0.01 |
|  | *T_30_* | 3238$\pm$258 | 7.38$\pm$0.03 | 45$\pm$3.3 | 1969$\pm$3.9 | 78$\pm$6.3 | 0.8$\pm$0.06 |
|  | *T_60_* | 3278$\pm$90 | 7.37$\pm$0.01 | 45$\pm$1.0 | 1982$\pm$3.6 | 78$\pm$2.1 | 0.8$\pm$0.02 |
|  | *T_95_* | 3169$\pm$80 | 7.39$\pm$0.01 | 49$\pm$0.9 | 2009$\pm$5.7 | 75$\pm$1.8 | 0.8$\pm$0.01 |

**Table S3**. Linear model results for *Siderastrea siderea* and *Pseudodiploria strigosa* host (calcification rate, total protein, total carbohydrates) and symbiont physiology (cell density and chlorophyll *a* concentration). Explicit models were determined from forward model selection, and are listed next to each species for the metric being considered. Sum Sq = sum of squares, Mean Sq = mean square of the error, NumDF = numerator degrees of freedom, and DenDF = denominator degrees of freedom.

| Factor | Sum Sq | Mean Sq | NumDF | DenDF | F-value | P-value |
| --- | --- | --- | --- | --- | --- | --- |
| CALCIFICATION |  | | | | | |
| *S. siderea* | Model = (calcification ~ *p*CO_2_ + duration + duration**p*CO_2_ + 1\|genotype)) | | | | | |
| duration | 4.9937 | 2.4968 | 2 | 189.99 | 10.6553 | <0.0001 |
| *p*CO_2_ | 20.0495 | 10.0247 | 2 | 190.02 | 42.7805 | <0.0001 |
| *p*CO_2_ *duration | 3.4102 | 0.8526 | 4 | 189.97 | 3.6382 | 0.0070 |
| *P. strigosa* | Model = (calcification ~ duration + temperature + *p*CO_2_ + reef zone + duration*temperature + temperature**p*CO_2_ + temperature*reef zone + 1\|genotype) | | | | | |
| duration | 20.0304 | 10.0152 | 2 | 158.13 | 27.2262 | <0.0001 |
| temperature | 17.8011 | 17.8011 | 1 | 159.19 | 48.392 | <0.0001 |
| *p*CO_2_ | 6.6279 | 3.3139 | 2 | 158.49 | 9.0089 | 0.0002 |
| reef zone | 4.46 | 4.46 | 1 | 3.19 | 12.1243 | 0.036 |
| duration:temperature | 9.0137 | 4.5068 | 2 | 158.13 | 12.2518 | <0.0001 |
| temperature:*p*CO_2_ | 5.3588 | 2.6794 | 2 | 158.49 | 7.284 | 0.0009 |
| temperature:reef zone | 1.7523 | 1.7523 | 1 | 159.37 | 4.7635 | 0.031 |
| TOTAL PROTEIN |  | | | | | |
| *S. siderea* | Model = (protein ~ duration + temperature + 1\|genotype) | | | | | |
| duration | 1.29E+00 | 4.29E-01 | 3 | 133 | 10.0511 | <0.0001 |
| temperature | 3.03E-01 | 3.03E-01 | 1 | 133 | 7.0908 | <0.01 |
| *P. strigosa* | Model = (protein ~ temperature + 1\|genotype) | | | | | |
| temperature | 1.48E+00 | 1.48E+00 | 1 | 111.08 | 28.712 | <0.0001 |
| CARBOHYDRATES |  | | | | | |
| *S. siderea* | Model = (carbohydrate ~ temperature + 1\|genotype) | | | | | |
| temperature | 2.47E+01 | 2.47E+01 | 1 | 135 | 9.146 | <0.01 |
| *P. strigosa* | Model = (carbohydrate ~ temperature + duration + duration*temperature + 1\|genotype) | | | | | |
| temperature | 4.96E+01 | 4.96E+01 | 1 | 102.22 | 25.9934 | <0.0001 |
| duration | 3.70E+01 | 1.23E+01 | 3 | 102.1 | 6.4564 | <0.001 |
| temperature:duration | 3.02E+01 | 1.01E+01 | 3 | 102.02 | 5.2671 | 0.002 |
| SYMBIONT DENSITY |  | | | | | |
| *S. siderea* | Model = (symbiont density ~ duration + temperature + *p*CO_2_ + duration**p*CO_2_ + 1\|genotype) | | | | | |
| duration | 5.01E+13 | 1.67E+13 | 3 | 301105 | 6.5304 | <0.0001 |
| temperature | 1.50E+13 | 1.50E+13 | 1 | 2478940 | 5.8538 | <0.01 |
| *p*CO_2_ | 1.62E+13 | 8.12E+12 | 2 | 701697 | 3.1759 | 0.04 |
| duration: *p*CO_2_ | 4.75E+13 | 7.92E+12 | 6 | 578830 | 3.0962 | 0.005 |
| *P. strigosa* | Model = (symbiont density ~ temperature + duration + 1\|genotype) | | | | | |
| duration | 4.64E+13 | 4.64E+13 | 1 | 696674 | 19.554 | <0.0001 |
| temperature | 4.81E+13 | 1.60E+13 | 3 | 142806 | 6.7636 | <0.001 |
| CHLOROPHYLL A |  | | | | | |
| *S. siderea* | Model = (Chl *a* ~ duration + *p*CO_2_ + 1\|genotype) | | | | | |
| duration | 1.57E+03 | 5.22E+02 | 3 | 127.36 | 40.7989 | <0.0001 |
| *p*CO_2_ | 1.75E+02 | 8.77E+01 | 2 | 127.32 | 6.8468 | <0.001 |
| *P. strigosa* | Model = (Chl *a* ~ temperature + *p*CO_2_ + (1\|genotype) | | | | | |
| temperature | 1.26E+03 | 1.26E+03 | 1 | 108.22 | 35.9079 | <0.0001 |
| *p*CO_2_ | 2.61E+02 | 1.30E+02 | 2 | 108.05 | 3.7166 | 0.028 |

**Table S4.** Summary of Tukey’s HSD post-hoc tests for *Siderastrea siderea* and *Pseudodiploria strigosa* host (calcification, total protein, total carbohydrates) and symbiont physiology (cell density and chlorophyll *a* concentration). Only significant interactions are included. SE = standard error and DF = degrees of freedom.

| **Contrast** | **Estimate** | **SE** | **DF** | **T-ratio** | **P-value** |
| --- | --- | --- | --- | --- | --- |
| CALCIFICATION | | | | | |
| ***Siderastrea siderea*** | | | | | |
|  | Comparison = *p*CO_2_ | | | | |
| present day - extreme | 0.48 | 0.115 | 158 | 4.186 | 0.0001 |
| next century - extreme | 0.316 | 0.114 | 159 | 2.765 | 0.0174 |
|  | Comparison = *p*CO_2_ \| duration | | | | |
| present day,T30 - extreme,T30 | 0.4803 | 0.115 | 158 | 4.186 | 0.0015 |
| present day,T30 - present day,T60 | 0.6926 | 0.104 | 158 | 6.691 | <.0001 |
| present day,T30 - present day,T95 | 0.658 | 0.13 | 158 | 5.056 | <.0001 |
| next century,T30 - next century,T60 | 0.6926 | 0.104 | 158 | 6.691 | <.0001 |
| next century,T30 - next century,T95 | 0.658 | 0.13 | 158 | 5.056 | <.0001 |
| extreme,T30 - extreme,T60 | 0.6926 | 0.104 | 158 | 6.691 | <.0001 |
| extreme,T30 - extreme,T95 | 0.658 | 0.13 | 158 | 5.056 | <.0001 |
|  | Comparison = reef zone \| duration \| *p*CO_2_\| temperature | | | | |
| duration = T60, pco2 = extreme, temperature = 28; FR -- NS | 0.6159 | 0.2702 | 2.279 | 68 | 0.0258 |
| duration = T95, pco2 = present day, temperature = 31; FR -- NS | 0.7916 | 0.3627 | 2.183 | 68 | 0.0325 |
| duration = T95, pco2 = next century, temperature = 31; FR -- NS | -1.0057 | 0.3627 | -2.773 | 68 | 0.00716 |
| duration = T60, pco2 = extreme, temperature = 31; FR -- NS | 0.9449 | 0.2811 | 3.362 | 68 | 0.00127 |
| ***Pseudodiploria strigosa*** | | | | | |
|  | Comparison = temperature | | | | |
| 28 - 31 | 0.734 | 0.106 | 159 | 6.939 | <.0001 |
|  | Comparison = reef zone | | | | |
| FR - NS | -0.511 | 0.147 | 3.08 | -3.477 | 0.0385 |
|  | Comparison = pCO2 | | | | |
| present day - next century | 0.165 | 0.112 | 158 | 1.472 | 0.3071 |
| present day - extreme | 0.48 | 0.115 | 158 | 4.186 | 0.0001 |
| next century - extreme | 0.316 | 0.114 | 159 | 2.765 | 0.0174 |
|  | Comparison = temperature \| duration | | | | |
| 28,T30 - 28,T60 | 0.5579 | 0.15 | 158 | 3.726 | 0.0036 |
| 31,T30 - 31,T60 | 0.8272 | 0.143 | 158 | 5.786 | <.0001 |
| 31,T30 - 31,T95 | 1.3013 | 0.181 | 158 | 7.196 | <.0001 |
| 28,T60 - 31,T60 | 0.4846 | 0.164 | 159 | 2.958 | 0.0409 |
| 28,T60 - 28,T95 | -0.5432 | 0.2 | 158 | -2.718 | 0.0773 |
| 28,T60 - 31,T95 | 0.9587 | 0.198 | 159 | 4.848 | <.0001 |
| 31,T60 - 28,T95 | -1.0278 | 0.198 | 158 | -5.186 | <.0001 |
| 28,T95 - 31,T95 | 1.5019 | 0.227 | 158 | 6.616 | <.0001 |
|  | Comparison = temperature \| reef zone | | | | |
| 28,F - 31,F | 0.944 | 0.16 | 159.8 | 5.914 | <.0001 |
| 31,F - 31,N | -0.72 | 0.171 | 5.72 | -4.21 | 0.0238 |
|  | Comparison = temperature \| *p*CO_2_ | | | | |
| 28,present day - 31,present day | 1.1908 | 0.167 | 158 | 7.137 | <.0001 |
| 28,present day - 31,next century | 1.105 | 0.167 | 158 | 6.623 | <.0001 |
| 28,present day - 28,extreme | 0.9152 | 0.168 | 159 | 5.457 | <.0001 |
| 28,present day - 31,extreme | 1.2362 | 0.167 | 158 | 7.409 | <.0001 |
| 31,present day - 28,next century | -0.7758 | 0.164 | 158 | -4.741 | 0.0001 |
| 28,next century - 31,next century | 0.69 | 0.164 | 158 | 4.216 | 0.0006 |
| 28,next century - 28,extreme | 0.5002 | 0.166 | 159 | 3.009 | 0.0354 |
| 28,next century - 31,extreme | 0.8212 | 0.164 | 158 | 5.018 | <.0001 |
|  | Comparison = reef zone \| duration \| *p*CO_2_\| temperature | | | | |
| duration = T30, pco2 = present day, temperature = 31; FR -- NS | -1.0357 | 0.3411 | -3.036 | 96 | 0.00308 |
| duration = T60, pco2 = present day, temperature = 31; FR -- NS | -0.8060 | 0.4105 | -1.963 | 96 | 0.05 |
| duration = T95, pco2 = next century, temperature = 31; FR -- NS | -1.3058 | 0.5701 | -2.291 | 96 | 0.0242 |
| duration = T30, pco2 = extreme, temperature = 31; FR -- NS | -0.8395 | 0.3411 | -2.461 | 96 | 0.0156 |
| PROTEIN | | | | | |
| ***Siderastrea siderea*** | | | | | |
|  | Comparison = temperature | | | | |
| 28 - 31 | 0.0937 | 0.0352 | 128 | 2.662 | 0.0088 |
|  | Comparison = duration | | | | |
| T0 - T60 | -0.22823 | 0.0512 | 129 | -4.462 | 0.0001 |
| T0 - T95 | -0.13566 | 0.0487 | 128 | -2.785 | 0.031 |
| T30 - T60 | -0.23759 | 0.0511 | 129 | -4.645 | <.0001 |
| T30 - T95 | -0.14502 | 0.0487 | 128 | -2.976 | 0.0181 |
| ***Pseudodiploria strigosa*** | | | | | |
|  | Comparison = temperature | | | | |
| 28 - 31 | 0.225 | 0.042 | 111 | 5.357 | <.0001 |
| CARBOHYDRATES | | | | | |
| ***Siderastrea siderea*** | | | | | |
|  | Comparison = temperature | | | | |
| 28 - 31 | 0.849 | 0.281 | 130 | 3.023 | 0.003 |
| ***Pseudodiploria strigosa*** | | | | | |
|  | Comparison = temperature | | | | |
| 28 - 31 | 1.33 | 0.261 | 102 | 5.095 | <.0001 |
|  | Comparison = duration | | | | |
| T0 - T60 | 1.377 | 0.375 | 102 | 3.67 | 0.0022 |
| T30 - T60 | 0.881 | 0.376 | 102 | 2.342 | 0.0953 |
| T60 - T95 | -1.5 | 0.376 | 102 | -3.989 | 0.0007 |
|  | Comparison = duration \| temperature | | | | |
| T0,28 - T60,31 | 2.209 | 0.497 | 102 | 4.446 | 0.0006 |
| T0,28 - T95,31 | 1.827 | 0.497 | 102 | 3.678 | 0.0087 |
| T30,28 - T95,28 | -1.881 | 0.522 | 102 | -3.602 | 0.0111 |
| T60,28 - T95,28 | -2.618 | 0.558 | 102 | -4.695 | 0.0002 |
| T95,28 - T0,31 | 2.073 | 0.524 | 102 | 3.957 | 0.0034 |
| T95,28 - T30,31 | 2.451 | 0.514 | 102 | 4.77 | 0.0002 |
| T95,28 - T60,31 | 3.475 | 0.514 | 102 | 6.764 | <.0001 |
| T95,28 - T95,31 | 3.094 | 0.514 | 102 | 6.022 | <.0001 |
| SYMBIONT DENSITY | | | | | |
| ***Siderastrea siderea*** | | | | | |
|  | Comparison = duration | | | | |
| T0 - T60 | 961879 | 398838 | 118 | 2.412 | 0.0804 |
| T0 - T95 | 1661877 | 379838 | 117 | 4.375 | 0.0002 |
|  | Comparison = temperature | | | | |
| 28 - 31 | 669132 | 276720 | 117 | 2.418 | 0.0171 |
|  | Comparison = pCO2 | | | | |
| present day - extreme | 819314 | 336723 | 117 | 2.433 | 0.0432 |
|  | Comparison = duration \| *p*CO_2_ | | | | |
| T30,present day - T95,present day | 2740085 | 653760 | 118 | 4.191 | 0.003 |
| ***Pseudodiploria strigosa*** | | | | | |
|  | Comparison = duration | | | | |
| T0 - T30 | 1178736 | 402554 | 106 | 2.928 | 0.0214 |
| T0 - T60 | 1357099 | 419508 | 107 | 3.235 | 0.0087 |
| T0 - T95 | 1731491 | 406231 | 106 | 4.262 | 0.0003 |
|  | Comparison = temperature | | | | |
| 28 - 31 | 1280321 | 289879 | 106 | 4.417 | <.0001 |
| CHLOROPHYLL A | | | | | |
| ***Siderastrea siderea*** | | | | | |
|  | Comparison = duration | | | | |
| T0 - T60 | -6.972 | 0.878 | 127 | -7.94 | <.0001 |
| T0 - T95 | -7.653 | 0.843 | 127 | -9.075 | <.0001 |
| T30 - T60 | -5.584 | 0.884 | 127 | -6.317 | <.0001 |
| T30 - T95 | -6.266 | 0.85 | 127 | -7.37 | <.0001 |
|  | Comparison = *p*CO_2_ | | | | |
| present day - extreme | 2.603 | 0.751 | 127 | 3.465 | 0.0021 |
| next century - extreme | 2.09 | 0.738 | 127 | 2.831 | 0.0148 |
| ***Pseudodiploria strigosa*** | | | | | |
|  | Comparison = *p*CO_2_ | | | | |
| present day - extreme | 3.7 | 1.36 | 108 | 2.721 | 0.0206 |
|  | Comparison = temperature | | | | |
| 28 - 31 | 6.6 | 1.1 | 109 | 5.985 | <.0001 |

**Table S5.** PCA Adonis summaries associated with Figure 5 and Figure S2. The Adonis model used to determine results is listed next to each species and time point. All Adonis models were run with 10,000 permutations. Significant p-values are bolded. Sum Sq = sum of squares, Mean Sq = mean square of the error, and DF = degrees of freedom.

| **Factor** | **DF** | **Sum Sq** | **Mean Sq** | **F-Model** | **R^2^** | **P-value** |
| --- | --- | --- | --- | --- | --- | --- |
| ***S. siderea,* T_30_** | Model = Adonis(scores ~ reef zone* *p*CO_2_*temperature*genotype) | | | | | |
| Reef zone | 1 | 0.000682 | 0.0006821 | 0.22531 | 0.00726 | 0.86741 |
| *p*CO_2_ | 2 | 0.016761 | 0.0083804 | 2.76814 | 0.1785 | 0.05369 |
| Temperature | 1 | 0.007265 | 0.0072649 | 2.39968 | 0.07737 | 0.11039 |
| Genotype | 4 | 0.013071 | 0.0032677 | 1.07936 | 0.13921 | 0.39876 |
| Reef zone: *p*CO_2_ | 2 | 0.004736 | 0.0023679 | 0.78215 | 0.05044 | 0.52045 |
| Reef zone:temperature | 1 | 0.000647 | 0.0006469 | 0.21368 | 0.00689 | 0.87991 |
| *p*CO_2_:temperature | 2 | 0.002629 | 0.0013143 | 0.43413 | 0.028 | 0.77862 |
| Reef zone: *p*CO_2_:temperature | 2 | 0.002694 | 0.0013471 | 0.44495 | 0.02869 | 0.76072 |
| ***S. siderea,* T_60_** | Model = Adonis(scores ~ reef zone* *p*CO_2_*temperature*genotype) | | | | | |
| Reef zone | 1 | 0.00015 | 0.0001505 | 0.1082 | 0.00335 | 0.9629 |
| *p*CO_2_ | 2 | 0.002017 | 0.0010086 | 0.7255 | 0.04486 | 0.52395 |
| Temperature | 1 | 0.004498 | 0.0044982 | 3.2356 | 0.10004 | 0.07389 |
| Genotype | 4 | 0.011229 | 0.0028073 | 2.0193 | 0.24973 | 0.12489 |
| Reef zone: *p*CO_2_ | 2 | 0.005664 | 0.0028321 | 2.0371 | 0.12597 | 0.14439 |
| Reef zone:temperature | 1 | 0.000524 | 0.0005243 | 0.3771 | 0.01166 | 0.66283 |
| *p*CO_2_:temperature | 2 | 0.002126 | 0.0010631 | 0.7647 | 0.04729 | 0.51335 |
| Reef zone: *p*CO_2_:temperature | 2 | 0.003463 | 0.0017315 | 1.2455 | 0.07702 | 0.30177 |
| ***S. siderea,* T_95_** | Model = Adonis(scores ~ reef zone* *p*CO_2_*temperature*genotype) | | | | | |
| Reef zone | 1 | 0.000763 | 0.00076323 | 1.4642 | 0.02308 | 0.21468 |
| *p*CO_2_ | 2 | 0.00545 | 0.00272503 | 5.2277 | 0.16485 | **0.0016** |
| Temperature | 1 | 0.001076 | 0.00107617 | 2.0645 | 0.03255 | 0.12469 |
| Genotype | 4 | 0.007173 | 0.00179329 | 3.4403 | 0.21696 | **0.0036** |
| Reef zone: *p*CO_2_ | 2 | 0.002143 | 0.00107126 | 2.0551 | 0.0648 | 0.09999 |
| Reef zone:temperature | 1 | 0.000569 | 0.000569 | 1.0916 | 0.01721 | 0.31037 |
| *p*CO_2_:temperature | 2 | 0.005689 | 0.0028447 | 5.4573 | 0.17208 | **0.0011** |
| Reef zone: *p*CO_2_:temperature | 2 | 0.000294 | 0.00014706 | 0.2821 | 0.0089 | 0.9527 |
| ***P. strigosa,* T_30_** | Model = Adonis(scores ~ reef zone* *p*CO_2_*temperature*genotype) | | | | | |
| Reef zone | 1 | 0.004577 | 0.0045768 | 1.10104 | 0.04311 | 0.3136 |
| *p*CO_2_ | 2 | 0.009332 | 0.0046661 | 1.12252 | 0.08791 | 0.3487 |
| Temperature | 1 | 0.008843 | 0.0088435 | 2.12747 | 0.0833 | 0.1493 |
| Genotype | 3 | 0.011136 | 0.0037121 | 0.89302 | 0.1049 | 0.4922 |
| Reef zone: *p*CO_2_ | 2 | 0.004723 | 0.0023613 | 0.56806 | 0.04449 | 0.6206 |
| Reef zone:temperature | 1 | 0.007089 | 0.007089 | 1.7054 | 0.06678 | 0.2053 |
| *p*CO_2_:temperature | 2 | 0.003006 | 0.001503 | 0.36159 | 0.02832 | 0.7722 |
| Reef zone: *p*CO_2_:temperature | 2 | 0.003417 | 0.0017085 | 0.41101 | 0.03219 | 0.7385 |
| ***P. strigosa,* T_60_** | Model = Adonis(scores ~ reef zone* *p*CO_2_*temperature*genotype) | | | | | |
| Reef zone | 1 | 0.015454 | 0.0154544 | 4.0395 | 0.09332 | 0.06189 |
| *p*CO_2_ | 2 | 0.035511 | 0.0177553 | 4.6409 | 0.21443 | **0.0293** |
| Temperature | 1 | 0.012789 | 0.0127888 | 3.3427 | 0.07722 | 0.08299 |
| Genotype | 3 | 0.037493 | 0.0124978 | 3.2667 | 0.2264 | 0.05309 |
| Reef zone: *p*CO_2_ | 2 | 0.01154 | 0.0057699 | 1.5082 | 0.06968 | 0.25787 |
| Reef zone:temperature | 1 | 0.009631 | 0.0096314 | 2.5175 | 0.05816 | 0.13939 |
| *p*CO_2_:temperature | 2 | 0.005624 | 0.0028118 | 0.735 | 0.03396 | 0.52145 |
| Reef zone: *p*CO_2_:temperature | 1 | 0.003129 | 0.0031294 | 0.818 | 0.0189 | 0.39766 |
| ***P. strigosa,* T_95_** | Model = Adonis(scores ~ reef zone* *p*CO_2_*temperature*genotype) | | | | | |
| Reef zone | 1 | 0.012884 | 0.0128841 | 2.299 | 0.07904 | 0.14029 |
| *p*CO_2_ | 2 | 0.002698 | 0.0013492 | 0.2408 | 0.01655 | 0.86961 |
| Temperature | 1 | 0.023802 | 0.0238021 | 4.2472 | 0.14602 | **0.0453** |
| Genotype | 3 | 0.011581 | 0.0038604 | 0.6888 | 0.07105 | 0.60454 |
| Reef zone: *p*CO_2_ | 2 | 0.005849 | 0.0029244 | 0.5218 | 0.03588 | 0.64354 |
| Reef zone:temperature | 1 | 0.022229 | 0.0222286 | 3.9665 | 0.13637 | 0.05349 |
| *p*CO_2_:temperature | 2 | 0.007571 | 0.0037856 | 0.6755 | 0.04645 | 0.55454 |
| Reef zone: *p*CO_2_:temperature | 2 | 0.003537 | 0.0017685 | 0.3156 | 0.0217 | 0.78892 |
| **Combined Species, T_30_** | Model = Adonis(scores ~ species * reef zone * *p*CO_2_ * temperature) | | | | | |
| Species | 1 | 0.054855 | 0.054855 | 14.4073 | 0.20515 | **0.0002** |
| *p*CO_2_ | 2 | 0.023192 | 0.011596 | 3.0456 | 0.08674 | **0.0472** |
| Temperature | 1 | 0.019842 | 0.019842 | 5.2114 | 0.07421 | **0.0233** |
| **Combined Species, T_60_** | Model = Adonis(scores ~ species * reef zone * *p*CO_2_ * temperature) | | | | | |
| Species | 1 | 0.054855 | 0.054855 | 14.4073 | 0.20515 | **0.0005999** |
| *p*CO_2_ | 2 | 0.023192 | 0.011596 | 3.0456 | 0.08674 | **0.0449955** |
| Temperature | 1 | 0.019842 | 0.019842 | 5.2114 | 0.07421 | **0.0186981** |
| **Combined Species, T_95_** | Model = Adonis(scores ~ species * reef zone * *p*CO_2_ * temperature) | | | | | |
| Species | 1 | 0.054855 | 0.054855 | 14.4073 | 0.20515 | **0.0004** |
| *p*CO_2_ | 2 | 0.023192 | 0.011596 | 3.0456 | 0.08674 | **0.046** |
| Temperature | 1 | 0.019842 | 0.019842 | 5.2114 | 0.07421 | **0.0222** |

**Table S6.** Linear model summaries associated with Figure 6 and Figure S4, including chi square values (Chi-Sq), degrees of freedom (DF) and significance (P-value) statistics. All data is associated with *P. strigosa,* and the table is separated by experimental time point (short-term = T_30_, moderate-term = T_60_, long-term = T_95_). Linear models are listed above the summaries for that model for reference.

| **Factor** | **Chi-Sq** | **DF** | **P-value** |
| --- | --- | --- | --- |
| ***P. strigosa, T_30_*** |  |  |  |
| PROTEIN | Model = (Calcification ~ Protein * Temperature + 1\|genotype) | | |
| Protein | 5.63 | 1 | 0.02 |
| Temperature | 0.12 | 1 | 0.73 |
| Protein:Temperature | 2.05 | 1 | 0.15 |
| CARBOHYDRATES | Model = (Calcification ~ Carbohydrates * Temperature + 1\|genotype) | | |
| Carbohydrates | 0.94 | 1 | 0.33 |
| Temperature | 0.04 | 1 | 0.83 |
| Carbohydrates:Temperature | 3.23 | 1 | 0.07 |
| SYMBIONT DENSITY | Model = (Calcification ~ Symbionts * Temperature + 1\|genotype) | | |
| Symbionts | 6.95 | 1 | 0.008 |
| Temperature | 0.45 | 1 | 0.50 |
| Symbionts:Temperature | 0.75 | 1 | 0.39 |
| ***P. strigosa, T_60_*** |  |  |  |
| PROTEIN | Model = (Calcification ~ Protein * Temperature + 1\|genotype) | | |
| Protein | 0.70 | 1 | 0.40 |
| Temperature | 3.83 | 1 | 0.05 |
| Protein:Temperature | 2.15 | 1 | 0.14 |
| CARBOHYDRATES | Model = (Calcification ~ Carbohydrates * Temperature + 1\|genotype) | | |
| Carbohydrates | 0.005 | 1 | 0.94 |
| Temperature | 4.53 | 1 | 0.03 |
| Carbs:Temperature | 5.72 | 1 | 0.017 |
| SYMBIONT DENSITY | Model = (Calcification ~ Symbionts * Temperature + 1\|genotype) | | |
| Symbionts | 0.18 | 1 | 0.67 |
| Temperature | 2.73 | 1 | 0.099 |
| Symbionts:Temperature | 5.76 | 1 | 0.016 |
| ***P. strigosa, T_95_*** |  |  |  |
| PROTEIN | Model = (Calcification ~ Protein * Temperature + 1\|genotype) | | |
| Protein | 5.09 | 1 | 0.024 |
| Temperature | 15.7 | 1 | < 0.0001 |
| Protein:Temperature | 4.06 | 1 | 0.044 |
| CARBOHYDRATES | Model = (Calcification ~ Carbohydrates * Temperature + 1\|genotype) | | |
| Carbohydrates | 0.74 | 1 | 0.39 |
| Temperature | 13.75 | 1 | < 0.001 |
| Carbs:Temperature | 3.44 | 1 | 0.064 |
| SYMBIONT DENSITY | Model = (Calcification ~ Symbionts * Temperature + 1\|genotype) | | |
| Symbionts | 13.3 | 1 | 0.0003 |
| Temperature | 15.76 | 1 | < 0.0001 |
| Symbionts:Temperature | 7.09 | 1 | 0.008 |
| CHLOROPHYLL A | Model = (Calcification ~ Chl *a ** Temperature + 1\|genotype) | | |
| Chl *a* | 13.5 | 1 | 0.0002 |
| Temperature | 4.63 | 1 | 0.03 |
| Chl *a:*Temperature | 1.45 | 1 | 0.22 |
